# How Glutamate Promotes Liquid-liquid Phase Separation and DNA Binding Cooperativity of *E. coli* SSB Protein

**DOI:** 10.1101/2022.01.17.476650

**Authors:** Alexander G. Kozlov, Xian Cheng, Hongshan Zhang, Min Kyung Shinn, Elizabeth Weiland, Binh Nguyen, Irina A. Shkel, Emily Zytkiewicz, Ilya J. Finkelstein, M. Thomas Record, Timothy M. Lohman

## Abstract

*E. coli* single-stranded-DNA binding protein (*Ec*SSB) displays nearest-neighbor (NN) and non-nearest-neighbor (NNN)) cooperativity in binding ssDNA during genome maintenance. NNN cooperativity requires the intrinsically-disordered linkers (IDL) of the C-terminal tails. Potassium glutamate (KGlu), the primary *E. coli* salt, promotes NNN-cooperativity, while KCl inhibits it. We find that KGlu promotes compaction of a single polymeric SSB-coated ssDNA beyond what occurs in KCl, indicating a link of compaction to NNN-cooperativity. *Ec*SSB also undergoes liquid-liquid phase separation (LLPS), inhibited by ssDNA binding. We find that LLPS, like NNN-cooperativity, is promoted by increasing [KGlu] in the physiological range, while increasing [KCl] and/or deletion of the IDL eliminate LLPS, indicating similar interactions in both processes. From quantitative determinations of interactions of KGlu and KCl with protein model compounds, we deduce that the opposing effects of KGlu and KCl on SSB LLPS and cooperativity arise from their opposite interactions with amide groups. KGlu interacts unfavorably with the backbone (especially Gly) and side chain amide groups of the IDL and therefore promotes amide-amide interactions in LLPS and NNN-cooperativity. By contrast, KCl interacts favorably with these amide groups and therefore inhibits LLPS and NNN-cooperativity. These results highlight the importance of salt interactions in regulating the propensity of proteins to undergo LLPS.

## Introduction

Single stranded (ss) DNA binding proteins (SSBs) are essential in all kingdoms of life. SSBs bind ssDNA intermediates formed transiently during genome maintenance to protect them from degradation and inhibit DNA secondary structures (Chase and Williams, 1986, Meyer and Laine, 1990, Lohman and Ferrari, 1994, Wold, 1997). *Escherichia coli* SSB (*Ec*SSB) also serves as a central hub for binding numerous metabolic proteins (SSB interacting proteins – SIPs) involved in replication, recombination and repair (Shereda et al., 2008).

*Ec*SSB functions as a homo-tetramer (Fig. 1A) (Lohman and Ferrari, 1994, Raghunathan et al., 2000), with each subunit (177 amino acids (aa)) composed of two domains (Fig. 1B): a structured N-terminal DNA binding domain (DBD) (residues 1-112), and a C-terminal domain (residues 113-177, Fig. 1D) composed of a flexible, intrinsically disordered linker (IDL) [56 aa] and a nine residue “acidic tip” (Fig. 1B). This acidic tip is conserved among many bacterial SSBs and is the primary site of interaction with the SIPs (Shereda et al., 2008, Genschel et al., 2000, Kozlov et al., 2010b, Marceau et al., 2011, Shereda et al., 2009, Ryzhikov and Korolev, 2012, Antony et al., 2013, Shinn et al., 2019). *Ec*SSB binds ssDNA in two major modes referred to as (SSB)_35_ and (SSB)_65_, where the subscripts denote the average number of ssDNA nucleotides occluded (Lohman and Overman, 1985, Bujalowski and Lohman, 1986). The relative stabilities of these modes depend on salt concentration and type, and protein to DNA ratio (binding density) (Chrysogelos and Griffith, 1982, Griffith et al., 1984, Lohman and Overman, 1985, Lohman et al., 1986, Bujalowski et al., 1988, Ferrari et al., 1994, Roy et al., 2007, Hamon et al., 2007, Kozlov et al., 2017, Kozlov et al., 2019), as well as applied force (Zhou et al., 2011, Suksombat et al., 2015, Hatch et al., 2008, Bell et al., 2015).

**Fig. 1.**
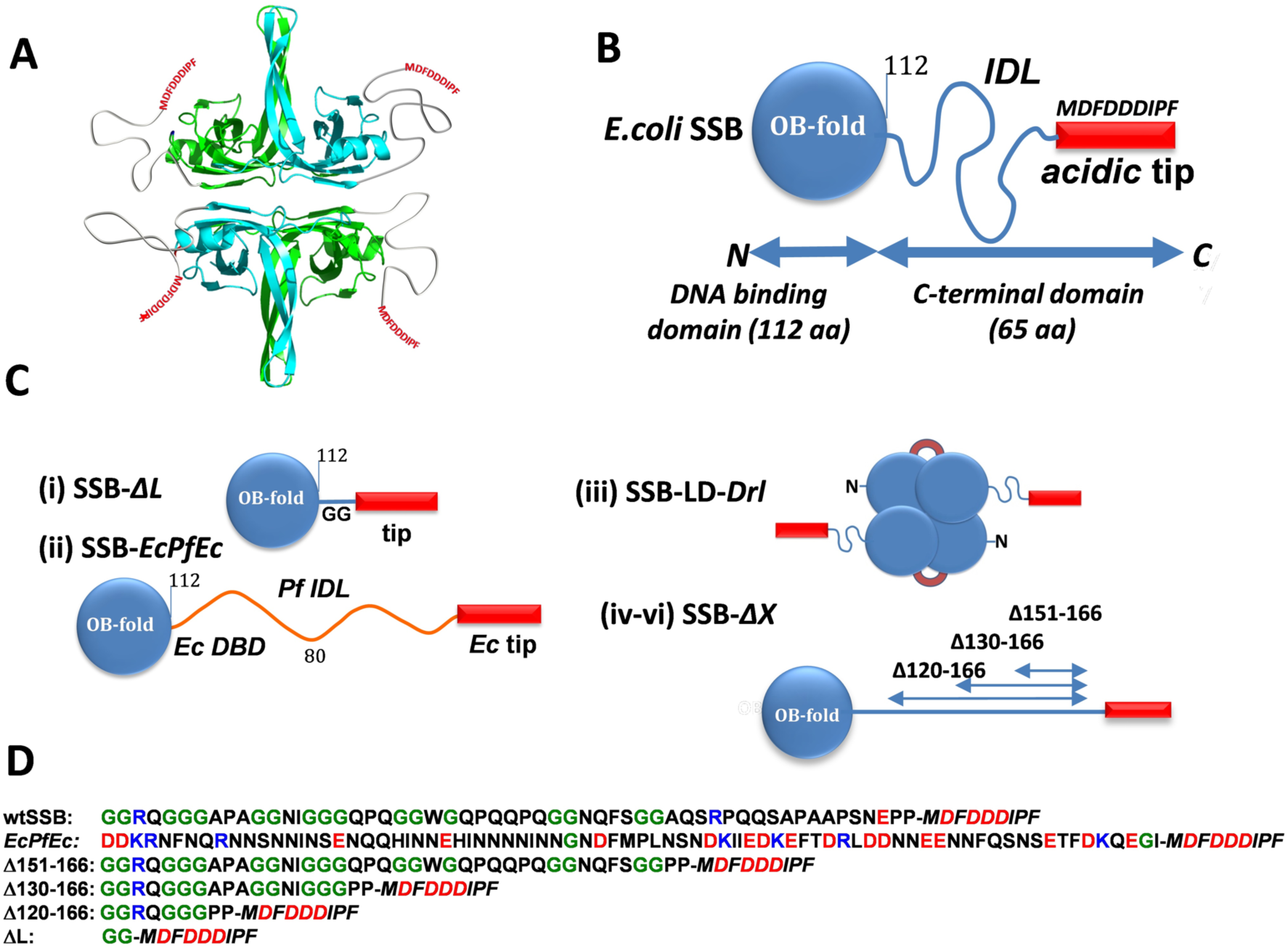
*E. coli* SSB constructs. **(A)** - cartoon of *Ec*SSB tetramer. Opposing subunits of the tetrameric core in cyan (front) and green (back) are shown with intrinsically disordered tails. **(B)** – schematic of *Ec*SSB subunit (177 aa), composed of an N-terminal DNA binding domain, OB fold (112 aa), intrinsically disordered linker, IDL (56 aa), and 9 aa conserved acidic tip (MDFDDDIPF). **(C)** – *Ec*SSB constructs: (i) – SSB-ΔL deletes the IDL (SSBΔ115-168); (ii) – SSB-*EcPfE*c chimera replaces the *Ec* IDL (56 aa) with the *Pf* IDL (80 aa); (iii) – Two-tailed *Ec*SSB dimeric construct (SSB-LD-*Drl*) in which 2 OB folds of upper and lower dimers are connected through *Deinococcus radiodurans* (*Dr*) linker; (iv-vi) – *Ec*SSB constructs containing different deletions within the IDL. **(D)** – Sequences of C-termini (intrinsically disordered IDL plus *Ec* tip (*Italic*)) for *Ec*SSB, *EcPfEc* chimera and all IDL deletion constructs; positively and negatively charged residues are shown in blue and red, respectively, glycines are shown in green.

In the (SSB)_65_ mode, favored at [NaCl]>0.20 M or [Mg^2+^]>10 mM at 25°C, the ssDNA wraps around all four subunits of the tetramer (Raghunathan et al., 2000) with a ∼65 nucleotide occluded site size. The topology of ssDNA wrapping in the (SSB)_65_ binding mode follows the seams on a baseball such that ssDNA enters and exits the tetramer in close proximity. On long ssDNA, the (SSB)_65_ mode displays "limited" cooperativity between adjacent tetramers (Chrysogelos and Griffith, 1982, Bujalowski and Lohman, 1987, Kozlov et al., 2019). In this mode SSB can diffuse along ssDNA destabilizing DNA hairpins and promoting RecA filament formation (Zhou et al., 2011, Roy et al., 2009).

In the (SSB)_35_ mode, favored at [NaCl]<10 mM or [MgCl_2_]<1 mM, and high SSB to DNA ratios (Lohman and Overman, 1985, Bujalowski and Lohman, 1986, Lohman et al., 1986), ssDNA interacts with only two subunits on average with a ∼35 nucleotide occluded site size. In this mode SSB binds ssDNA with unlimited nearest-neighbor (NN) cooperativity favoring formation of long protein clusters (Griffith et al., 1984, Lohman et al., 1986, Hamon et al., 2007, Sigal et al., 1972, Ferrari et al., 1994, Kozlov et al., 2015, Kozlov et al., 2019). Based on structural considerations it was suggested that SSB NN cooperativity might be promoted by interactions of adjacent tetramers through the L_45_ loops within the tetrameric core (Raghunathan et al., 2000) as well as directly through the residues of the core not involved in DNA binding (Dubiel et al., 2019, Kozlov et al., 2019). In this mode SSB can diffuse along ssDNA (Zhou et al., 2011, Suksombat et al., 2015) and undergo direct or intersegment transfer between separate ssDNA molecules (Kozlov and Lohman, 2002) or between distant sites on the same DNA molecule (Lee et al., 2014). The ability to undergo direct transfer appears to play a role in SSB recycling during replication (Kozlov and Lohman, 2002, Spenkelink et al., 2019).

Another level of non-nearest-neighbor (NNN) cooperativity has been identified recently for SSB bound to polymeric ssDNA (Bell et al., 2015, Kozlov et al., 2015, Kozlov et al., 2017). This NNN cooperativity occurs between SSB tetramers distantly bound to polymeric ssDNA and results in compaction/condensation of nucleoprotein complexes. Such interactions require the IDL (Kozlov et al., 2015, Kozlov et al., 2017) and are promoted by glutamate and acetate salts (Bell et al., 2015, Kozlov et al., 2015, Kozlov et al., 2017).

*Ec*SSB also undergoes liquid-liquid phase separation (LLPS) under solution conditions that mimic the *E. coli* environment. The intrinsically disordered C-terminal tails of SSB are essential for LLPS, which is suppressed by ssDNA (Harami et al., 2020). Here we explore the ability of *Ec*SSB to undergo LLPS as a function of temperature, salt type and concentration. We also explore how modifications within the IDL affect LLPS. We show that elevated concentrations of potassium glutamate (KGlu), the primary monovalent salt in *E. coli* (Richey et al., 1987, Record et al., 1998), promotes LLPS whereas KCl has the opposite effect. We present a thermodynamic analysis of interactions of KCl with protein model compounds and compare these with results for KGlu (Cheng et al., 2016) showing that these large opposing effects of KGlu and KCl on SSB LLPS likely result from their opposite interactions with backbone (especially G) and side chain amides of the SSB IDL in solution. We therefore propose that these amide groups interact with one another in the condensed phase, reducing or eliminating their interactions with water and salt ions. These amide-amide interactions, favored by KGlu and disfavored by KCl, appear to be important contributors to LLPS.

SSB LLPS, like NNN cooperative interactions of DNA-bound SSB, is a highly cooperative process. We find that conditions that promote LLPS also promote NNN cooperativity of SSB binding to ssDNA and conclude that similar cooperative interactions of tail residues drive these two processes mediated by ssDNA.

## Results

### Constructing phase diagrams using turbidity measurements

Liquid-liquid phase separation (LLPS) of *Ec*SSB, recently observed in the absence of ssDNA, is inhibited by binding of ssDNA (Harami et al., 2020). We refer to this phenomenon as LLPS, or simply phase separation (PS), although it has also been referred to as phase separation aided percolation (PSP) (Cohan and Pappu, 2020). LLPS of SSB is promoted by KGlu and requires the intrinsically disordered tails of SSB (Harami et al., 2020). Here, we explore how KGlu and KCl affect phase separation by determining their effect on the temperature corresponding to the phase boundary for different concentrations of SSB. These measurements were performed at different concentrations of KCl and KGlu using spectroscopic turbidity measurements (Amiram et al., 2011, Lyons et al., 2013, Cinar et al., 2019). The results of typical experiments at two SSB concentrations (3 and 12 µM) are shown in Fig. 2 (buffer T, 0.20 M KGlu). Starting at a temperature above the cloud point temperature (T_PS_) (25 °C and 50 °C, respectively), where the solutions are homogeneous, the temperature was decreased gradually. When the T_PS_ is reached (∼23 °C and 44 °C, respectively) the turbidity of the solution starts to increase. We use this as a proxy for the onset of phase separation. The further increase in turbidity reflects the growth of phase separated droplets (Harami et al., 2020). However, at some point this process slows and turbidity starts to decrease (12 µM, grey circles). Importantly, this process is reversible (3 µM, orange line) in the temperature range where turbidity rises, whereas reversibility is lost as the temperature is decreased further into the region where turbidity begins to decrease (12 µM, grey line). This irreversibility is supported by the fact that after the temperature reversal the SSB concentration decreased to 9 µM from the starting concentration of 12 µM, whereas the SSB concentration is maintained in the experiment performed at the lower [SSB] of 3 µM.

**Fig. 2.**
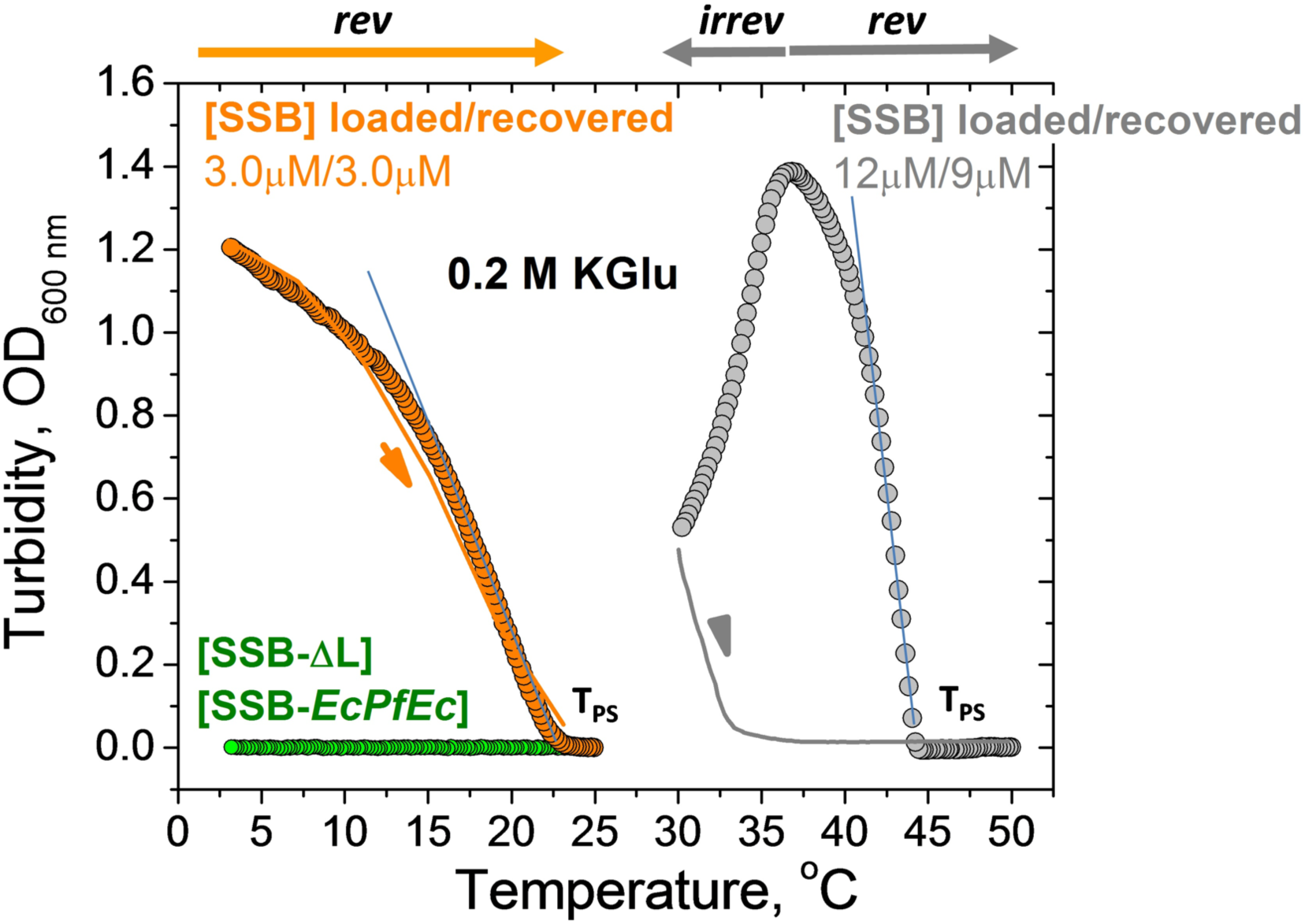
Determination of T_PS_ from turbidity measurements. Increase in turbidity (OD) monitored at 600 nm for wtSSB at 3 µM (orange circles) and 12 µM (gray circles) as the temperature decreases from 25°C to 3°C and from 50°C to 30°C, respectively, with ramp rate 0.2 °C/min (buffer T, 0.2M KGlu). T_PS_ in each case is obtained from extrapolation of the linear part of turbidity curve to the temperature axis. Corresponding reverse turbidity curves (orange and grey solid lines) obtained by increasing temperature with the rate 0.3 °C/min. No LLPS (change in turbidity) is observed for SSB-*EcPfEc* (13 µM) and SSB-ΔL (10 µM) from 25°C to 3°C (green circles, buffer T, 0.1M KGlu).

We used TIRF microscopy to monitor LLPS in the temperature range where the turbidity changes abruptly (see Fig. S1). An SSB solution (5 µM) containing 20 nM of SSB labeled with Cy5 in buffer T, 0.20 M KGlu prepared at 32 °C (above the T_PS_) was placed in a temperature controlled glass slide assembly. Rapid formation of dense phase droplets was observed as the temperature of the slide holder was maintained below T_PS_=30°C in the range where turbidity increases (see images taken at 27.5 °C in Fig. S1). However, the growth of freely diffusing droplets resulting in an increase in droplet size that is expected for a metastable phase (Rubinstein, 2003, Posey et al., 2018, Harami et al., 2020) was not observed. We suggest that droplet growth might be detectable at longer times. However, after 1-2 minutes the number of droplets decreases, presumably reflecting movement to the bottom of the slide channel under gravitational force and thus, disappearance from the focal plane of the microscope. Surprisingly, at temperatures near the maximum in turbidity we observe formation of fiber-like structures along with the freely diffusing liquid droplets (see images obtained for 22 °C in Fig. S1). These results show that turbidity measurements yield a reliable determination of the apparent T_PS_, at which the formation of dense phase starts to occur (Thomson et al., 1987, Petsev et al., 2003, Posey et al., 2018). We used this characteristic temperature to construct plots of T_PS_ vs [SSB] under different conditions. We use these as proxies for the low concentration arms of coexistence curves (Rubinstein, 2003, Posey et al., 2018) to examine the effects of salt concentration and type and SSB tail modifications on *Ec*SSB phase separation.

### Effects of glutamate vs. chloride on liquid-liquid phase separation of *Ec*SSB

We investigated the effects of KGlu and KCl concentrations on the phase behavior of wtSSB. Low concentration arms of coexistence curves (T_PS_ vs [SSB]) are shown in Fig. 3A for 6 concentrations of KGlu from 30 mM to 0.40 M. The upper limit on the concentration of SSB tetramers is ∼17 µM based on its solubility of 18-21 µM after dialysis at room temperature (22°C) in buffer T with no added salt. The upper temperature limit is ∼55 °C, above which the SSB tetramer begins to dissociate (Kozlov and Lohman, 1999). The coexistence curve in the [SSB]-T_PS_ plane has a parabolic shape (Fig. 3A) (Petsev et al., 2003, Rubinstein, 2003, Posey et al., 2018). We find that T_PS_ increases with increasing [KGlu] indicating that KGlu promotes phase separation and this effect is more pronounced at high SSB concentrations (Fig. 3A).

**Fig. 3.**
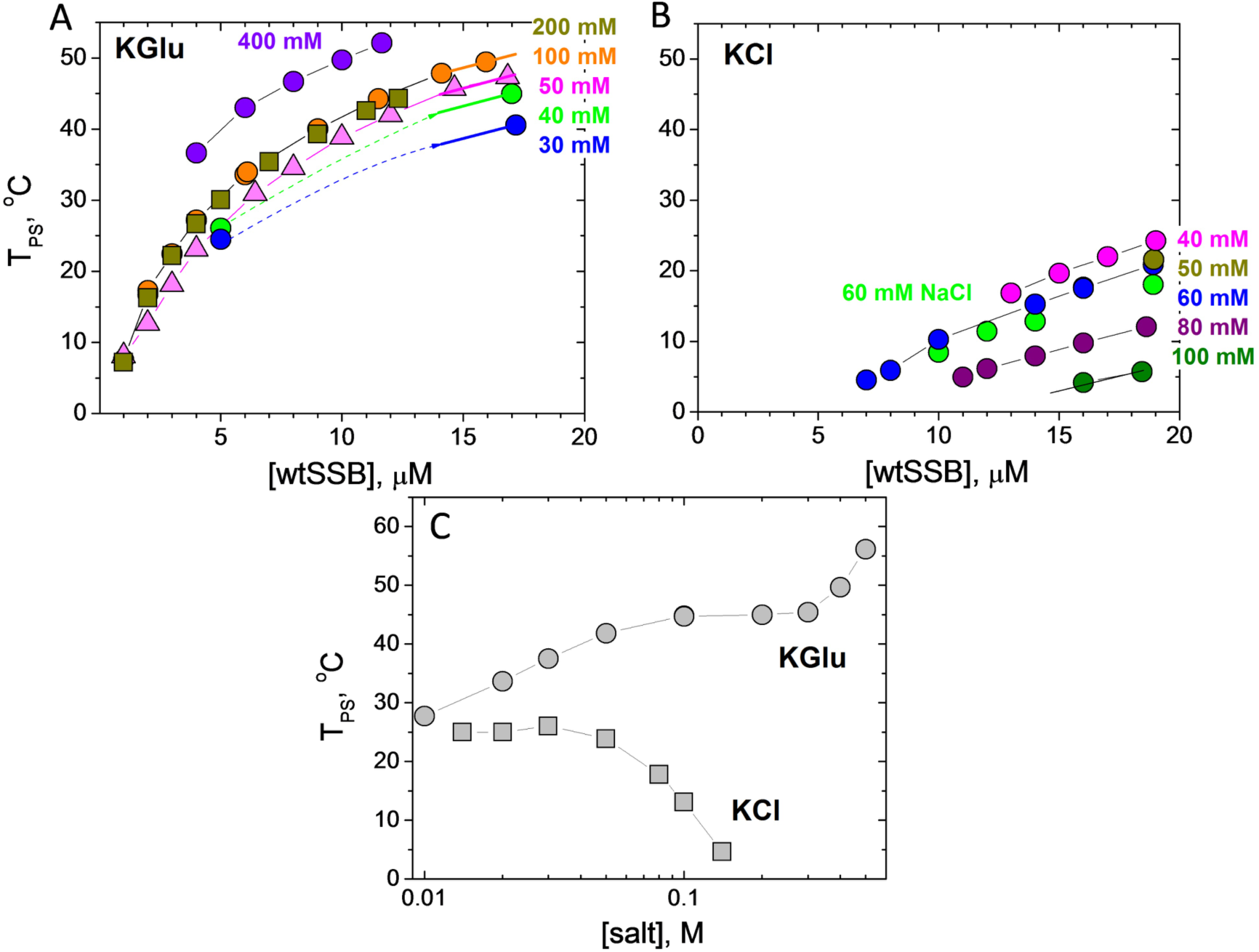
Increasing [Glu^-^] facilitates EcSSB LLPS, but increasing [Cl^-^] inhibits LLPS. SSB liquid-liquid phase diagrams (T_PS_ vs. [SSB], buffer T) obtained at different concentrations of **(A)** - KGlu: 30 mM (blue), 40 mM (green), 50 mM (magenta), 100 mM (orange), 200 mM (dark yellow) and 400 mM (violet); **(B)**- KCl: 40 mM (magenta), 50 mM (dark yellow), 60 mM (blue), 80 mM (purple), 100 mM (dark green) and 60 mM NaCl (green). **(C)** - SSB phase diagrams (T_PS_ vs [salt], buffer P) obtained for 8 µM SSB as a function of KCl (squares) and KGlu (circles).

In contrast, the effect of [KCl] on T_PS_ of SSB differs qualitatively from that of [KGlu] (Fig. 3B). In KCl, all T_PS_ values are shifted to lower temperatures and show the opposite dependence on [KCl]. The T_PS_ is nearly constant at low [KCl] (< 60 mM) but then decreases gradually, indicating that KCl disfavors phase separation, eventually eliminating condensate formation at high [KCl]. Effects of 60 mM KCl and NaCl (blue and green circles in Fig. 3B) are essentially the same indicating that interactions of K^+^ and Na^+^ with these SSB groups are the same (see Discussion). We note that the addition of 50 mM NaCl significantly decreases the ability of SSB to undergo phase separation in 0.10 M KGlu (compare orange and green circle dependences in Fig. 4A), although it does not eliminate phase separation (compare with Fig. 3B).

**Fig. 4.**
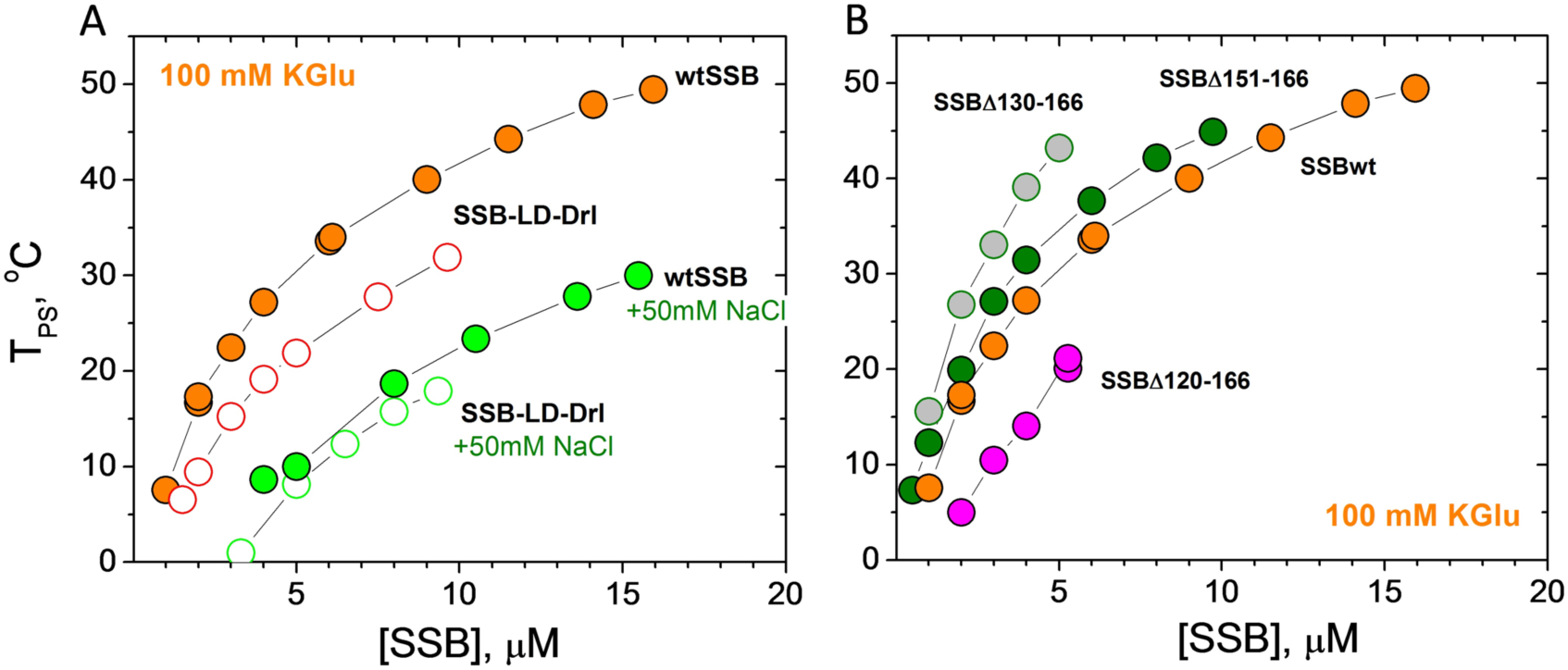
Number of tails and IDL deletions affect *Ec*SSB LLPS. SSB liquid-liquid phase diagrams (T_PS_ vs [SSB], buffer T, 0.1M KGu) obtained for: **(A)** - two tailed SSB-LD-*Drl* (open orange circles) and **(B)** - SSB with IDL deletions, Δ151-166 (green), Δ130-166 (grey), Δ120-166 (magenta). Phase diagrams of wtSSB for the same conditions are shown for comparison, including effect of the addition of 50 mM NaCl (panel A, open orange circles for wtSSB and open green circles for SSB-LD-*Drl*).

In the Tris buffer used to obtain the data in Fig. 3A and 3B, the pH drops by nearly a full pH unit from high to low temperature. Therefore, we also performed experiments in a phosphate buffer (Buffer-P, pH 7.5) for which the pH changes less dramatically from 7.61 to 7.46 in the range from 2 to 55 °C. These experiments were performed at constant [SSB] (8 µM tetramer), and the T_PS_ was determined as a function of [KCl] and [KGlu] (Fig. 3C). We observe a biphasic dependence of T_PS_ on [KGlu]. We observe an initial increase in T_PS_ that reaches a plateau between 0.10-0.30 M KGlu, followed by a further increase in T_PS_ above 0.40 M KGlu. In contrast, there is little effect of [KCl] until concentrations > 0.05 M, where increasing [KCl] inhibits phase separation. These dramatic differences between KCl and KGlu are due to differences in the interactions of Cl^-^ and Glu^-^ with SSB groups that are exposed in solution but buried in forming the SSB-SSB interactions that drive LLPS.

### Phase separation of *Ec*SSB is dependent on its intrinsically disordered linker (IDL)

Harami et al.(Harami et al., 2020) showed that removal of the intrinsically disordered *Ec*SSB tail eliminates phase separation of SSB under the solution conditions used for their experiments. In contrast, removal of the conserved acidic tip only weakens phase separation, but does not eliminate it. Here we show that the IDL of the C-terminal tail is necessary for phase separation even at low micromolar SSB concentrations. An SSB-ΔL construct, in which the IDL was deleted but the acidic tip remains (Δ113-168), does not undergo phase separation under conditions that promote it for wtSSB (0.10 M KGlu) within the temperature and concentration range examined (2 – 55 °C and 1 – 18 µM SSB, respectively, see Fig.2). The same is true for a chimeric construct, SSB-*EcPfEc* (Fig. 1C(ii)), in which the IDL of *E. coli* SSB was replaced with the longer and more highly charged IDL *of P. falciparum* SSB (Fig. 1D). This indicates that not only the presence but also the amino acid sequence/composition of the IDL is critical for phase separation.

We next examined how the number of SSB tails and the length of the IDL affects phase separation. We examined an SSB variant, SSB-LD-*Drl*, with only two tails as well as three variants with different length deletions within the IDL (SSB*Δ*151-166; SSB*Δ*130-166, SSB*Δ*120-166) (Fig. 1C and 1D). Figure 4A (open orange circles) shows that the two tailed variant is less effective at undergoing phase separation in 0.10 M KGlu. This indicates the importance of the multi-valency resulting from having four tails. Similar to wtSSB, addition of 50 mM NaCl significantly decreases T_PS_ of SSB-LD-*Drl* (Fig. 4A, open green circles). Interestingly, in these conditions (0.10 M KGlu + 50 mM NaCl) the coexistence curves of SSB-LD-*Drl* and wtSSB are essentially the same (Fig. 4A, open and filled green circles) indicating that the number of tails (two vs four) has a much smaller effect on phase separation at elevated [NaCl].

The low concentration arms of coexistence curves determined for the three SSB variants with partial deletions of the IDL are shown in Fig. 4B. Variants deleting 16 and 37 residues at the junction with the acidic tip (*Δ*151-166 and *Δ*130-166) increase T_PS_ and therefore promote phase separation compared to wtSSB. The effect of the 37-residue deletion is greater than that of the 16-residue deletion. However, deletion of an additional 10 residues (*Δ*120-166) reduces T_PS_ from that of wtSSB, indicating that this construct is less effective at undergoing phase separation (Fig. 4B). As mentioned above, the SSB-ΔL variant in which the entire linker is deleted (*Δ*113-168) does not undergo phase separation under the same solution conditions. That the shorter deletions, *Δ*151-166 and *Δ*130-166, promote phase separation was unexpected (see Discussion). However, we note that the propensity of these variants to still undergo phase separation correlates with their ability to form NNN cooperative complexes on polymeric M13 ssDNA (Kozlov et al., 2015, Kozlov et al., 2017).

We also find that a peptide corresponding to the entire *Ec*SSB tail (residues 113-117) does not undergo phase separation at the highest concentrations that we can achieve (600-800 µM) even at the lowest temperature examined (3 _o_C). Although this might be related to the fact that the effective concentration of the C-terminal tails of one wtSSB tetramer is calculated to be in the mM range, it is likely that the multi-valency of the tetrameric SSB core is required for LLPS. In addition, the SSB core has weak interactions with the negatively charged tips of the tails (Kozlov et al., 2010a, Su et al., 2014, Kozlov et al., 2015, Kozlov et al., 2017). In fact, we find that an SSB construct in which the four negative charges in the tip (Asp) are reversed to positive charges, *MKFKKKIPF* does not undergo phase separation in 0.10 M KGlu (data not shown) suggesting that the tip-core interactions are likely important for phase separation (see Discussion).

### DNA binding to SSB inhibits phase separation

Whereas wtSSB alone undergoes phase separation in 0.10 M KGlu as shown above, this phase separation is reversed upon SSB binding to ssDNA. Under conditions where we observe SSB droplets (4 µM), we note that droplets disappear upon addition of (dT)_35_ at a 1:1 stoichiometric ratio with wtSSB tetramer (Fig. S2B). Furthermore, solutions containing preformed 1:1 complexes of wtSSB with dT_35_ or dT_70_ ([SSB]=15 µM) do not show any indication of turbidity in 0.10 M KGlu even at the lowest temperature examined, 3 °C (Fig. S2A). These results indicate that phase separation is eliminated even when SSB is only partially bound with ssDNA in (SSB)_35_ mode (see Discussion). These observations are consistent with those of Harami et al. (Harami et al., 2020).

### Determinants of SSB phase separation correlate with the ability to form NNN complexes on long single stranded DNA

*Ec*SSB displays high non-nearest-neighbor (NNN) cooperativity on natural polymeric ssDNA (e.g., ss M13 phage DNA) at low [NaCl]/[KCl] as indicated by a bimodal distribution of SSB-ssDNA complexes at less than saturating SSB concentrations in gel electrophoresis and sedimentation velocity experiments (Lohman et al., 1986, Kozlov et al., 2015). This NNN cooperativity is also promoted at high [KGlu] and [acetate] (>0.10 M), but is inhibited at high [KCl] (Kozlov et al., 2017) (see Fig. S3). This cooperativity results from interactions between SSB tetramers distantly bound to ssDNA and requires the IDL (Kozlov et al., 2015).

The conditions that promote NNN cooperativity of *Ec*SSB binding to polymeric ssDNA are similar to those that promote SSB phase separation. To explore this further, we used ssDNA curtains on low complexity ssDNA (Schaub et al., 2018) to examine the SSB-induced collapse of ssDNA as a function of [KCl] and [KGlu] (Fig. 5A). For these experiments we used both wtSSB and the SSB-ΔL variant, which does not form NNN complexes on ssDNA. The low complexity ssDNA contains only thymine and cytosine bases and thus does not form any internal base pairs. One end of ssDNA is tethered to the surface, and the other end is fluorescently labeled via an anti dsDNA-antibody. By fluorescently tracking the free DNA end, we can monitor changes in the ssDNA length due to the addition of *Ec*SSB in different buffers (Fig. 5B). We first confirmed that changing the salt concentration and type does not influence the length of unbound ssDNA (Fig. S4). Next, we monitored the salt-dependent changes in ssDNA incubated with excess *Ec*SSB. *Ec*SSB was first added to the flow cells in imaging buffer without added salt. These conditions favor SSB binding in its (SSB)_35_ mode. All free *Ec*SSB was then washed out of the flow cell. The imaging buffer was then switched from 0 to 300 mM KCl, conditions that promote formation of the more compact, fully wrapped (SSB)_65_ complex. Hence the *Ec*SSB-ssDNA filament undergoes a dramatic compaction as shown previously (Schaub et al., 2018). However, when the 300 mM KCl is replaced with 300 mM KGlu a further compaction of the ssDNA occurs beyond that expected from formation of the (SSB)_65_ binding mode. This additional compaction is reversed when the imaging buffer is switched back to 300 mM KCl (Fig. 5C). We quantified the ssDNA length across multiple molecules and plotted the length in 300 mM KCl and 300 mM KGlu vs. the length in the absence of added salt (0 mM salt). These showed a linear fit with a slope of 0.53 for the 300 mM KCl data (N=29 ssDNA:*Ec*SSB molecules) and 0.42 for the 300 mM KGlu data (N=42 molecules) (Fig. 5D) indicating that the SSB-ssDNA was compacted more in the presence of KGlu compared to KCl (Fig. 5E).

**Fig. 5.**
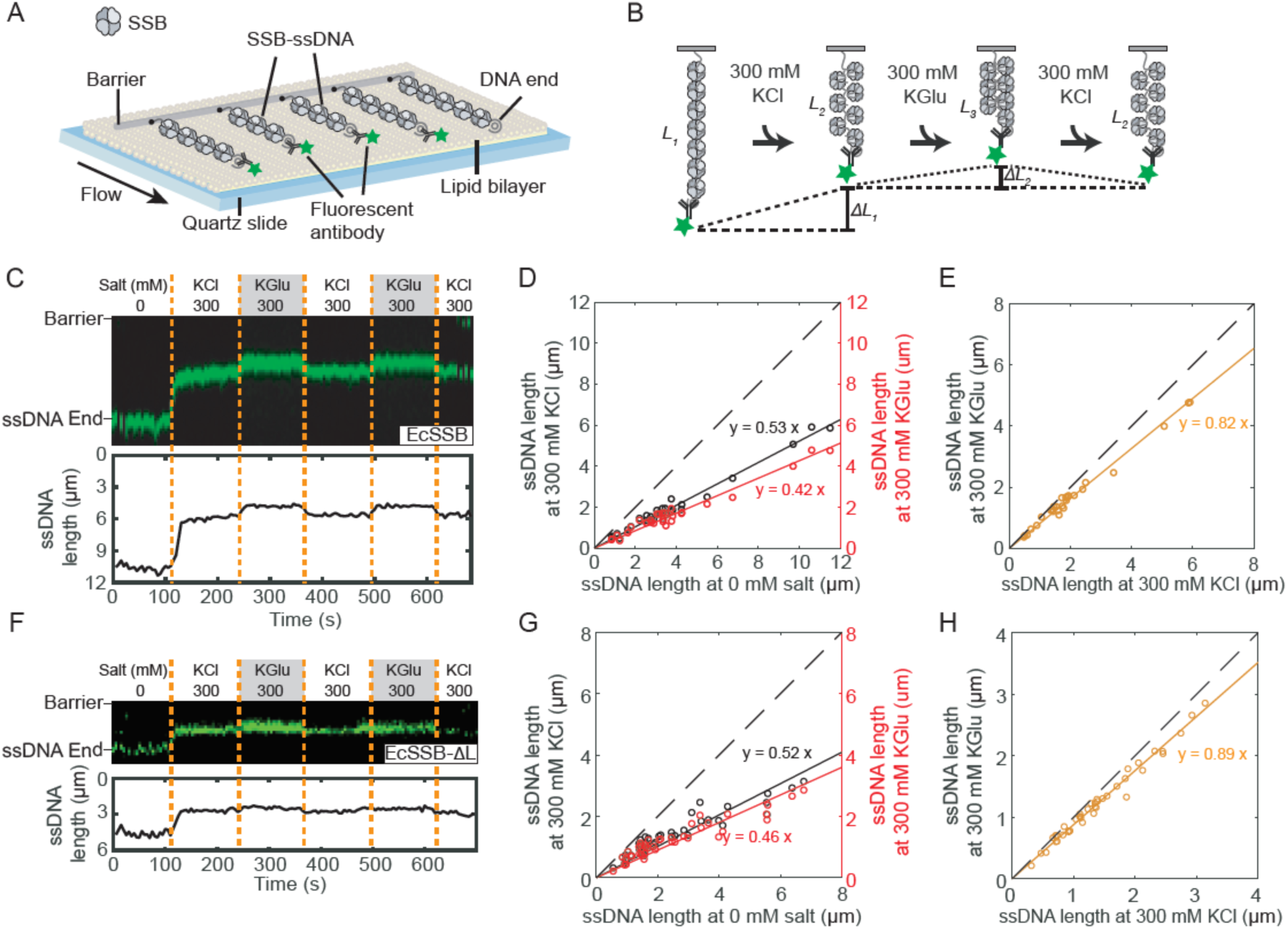
Glutamate promotes additional compaction of single polymeric DNA molecules coated with SSB. Illustration of ssDNA curtains decorated with SSB. (**B**) Schematic of the salt exchange assay. The length of SSB-ssDNA is changed when different buffer is loaded. (**C**) Representative kymograph (top) and single-particle tracking (bottom) showing the compaction of SSB-coated ssDNA end (green). Dashed orange lines denote when the buffer was switched. (**D**) Correlation between ssDNA-SSB lengths at 0 and 300 mM KCl (black), and at 0 and 300 mM KGlu (red). The solid lines are a linear fit to the data (N = 29 molecules). The dashed line represents a slope of 1. (**E**) Correlation between ssDNA-SSB lengths at 300 mM KCl and 300 mM KGlu (orange). (**F**) Representative kymograph and single-particle tracking showing the compaction of SSB-*ΔL*-coated ssDNA. (**G**) Correlation between length of ssDNA coated with *Ec*SSB-ΔL at 0 and 300 mM KCl, and at 0 and 300 mM KGlu (N = 42 molecules). (**H**) Correlation between length of ssDNA coated with SSB-*ΔL* at 300 mM KCl and 300 mM KGlu.

Experiments with SSB-ΔL showed a much smaller effect of KGlu, on ssDNA compaction (Figs. 5F, 5G, 5H). These experiments show that the large additional compaction of wtSSB-ssDNA complexes promoted by KGlu correlates with the promotion by KGlu of NNN cooperativity of wtSSB (Kozlov et al., 2015) and that the *Ec*SSB IDL is required for both effects. Furthermore, the conditions that promote SSB-DNA compaction and NNN cooperativity are the same as those that promote phase separation.

Interestingly, whereas Bell et al. (Bell et al., 2015) showed an additional compaction (condensation) of SSB-ssDNA complexes beyond that expected for formation of the fully wrapped (SSB)_65_ binding mode, Schaub et al. (Schaub et al., 2018), did not observe this additional SSB-DNA compaction. However, the experiments of Bell et al. (Bell et al., 2015) were performed in buffers containing sodium acetate as the added monovalent salt, whereas the Schaub et al. (Schaub et al., 2018) experiments were performed in buffers containing NaCl. This apparent discrepancy can be explained by the observation that, like glutamate salts, high acetate salt concentrations also promote NNN cooperativity, whereas chloride salts do not (Kozlov et al., 2017).

### Interactions of KCl with Model Compounds With the Most Common Functional Groups of Proteins; Comparison with KGlu

We hypothesize that the dramatic differences in the effects of KGlu vs. KCl on *Ec*SSB phase separation are due to the preferential interactions of these salts with the *Ec*SSB C-terminal tails. To assess this, here we determine the interactions of KCl with a series of model compounds displaying subsets of the functional groups of proteins. KGlu was the subject of an analogous previous study (Cheng et al., 2016).

Results of osmolality measurements on solutions of KCl and 18 model compounds and effects of KCl on the solubility of naphthalene are shown in Figure S5. Panels A-C plot osmolality differences *Δ*Osm (Eq. 1 in Methods) as a function of the product of model compound (m_2_) and KCl (m_3_) molalities. These plots are linear over the concentration ranges investigated and chemical potential derivatives (∂μ_2_/∂m_3_)_T.P,m2_ = μ_23_ (Eq. 2 in Methods), listed in Table S1, are obtained from the slopes. A negative μ_23_ indicates a favorable interaction; interactions of KCl with the most polar compounds studied (urea, formamide, malonamide; glycine, alanine) are favorable while μ_23_ becomes increasingly unfavorable as the amount of hydrocarbon in the model compound increases. Panel D of Fig S5 plots the logarithm of the solubility of naphthalene as a function of KCl concentration; the μ_23_ value quantifying the very unfavorable KCl-naphthalene interaction is obtained from the slope (see Eq. 3 of Methods).

Table S1 compares these μ_23_ values with those determined previously for KGlu (Cheng et al., 2016). For all compounds investigated, KGlu μ_23_ values are much less favorable/much more unfavorable than for KCl, in some cases by an order of magnitude. The only favorable interaction of KGlu is with glycine, explained as net-favorable interactions of the ions of KGlu with the ammonium and carboxylate groups of glycine.

Dissection of the set of KCl-model compound μ_23_ values using ASA information for these model compounds (Cheng et al., 2016, Cheng et al., 2017) yields *α*-values (Eq. 4 in Methods) that quantify the intrinsic strengths of interaction of KCl with unit areas of the 6 most common types of protein O, N and C unified atoms. Predicted μ_23_ values obtained from these *α*-values are compared with observed *α*-values in Fig. S6 and Table S1.

KCl *α*-values are compared with KGlu *α*-values in Fig. 6. The bar graph shows that both salts interact unfavorably with both aliphatic and aromatic hydrocarbon (sp^3^C and sp^2^C unified atoms), but exhibit opposite interactions with polar and charged N and O unified atoms. KCl interacts favorably with amide and carboxylate sp^2^O and unfavorably with amide sp^2^N and cationic N. By contrast, KGlu interacts unfavorably with amide and carboxylate sp^2^O and favorably with amide sp^2^N and cationic N.

**Fig. 6.**
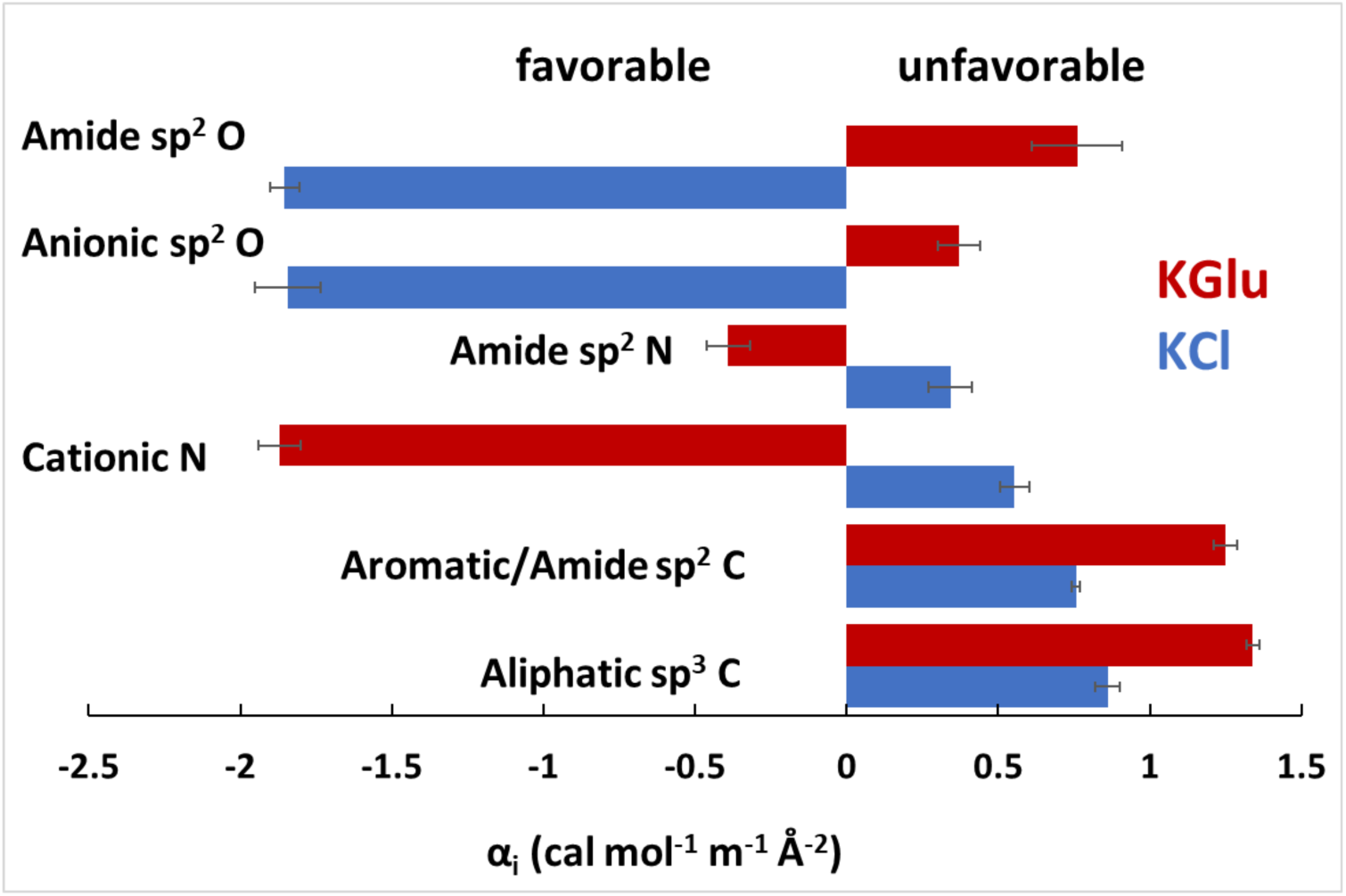
Intrinsic Strengths of Interaction of KGlu and KCl with Unified C, N and O Atoms of Proteins. Interaction potentials (α-values: Eq. 4 and Table S2) quantify the interactions of KCl (blue) and KGlu (Cheng et al., 2016) (red) with a unit area of each type of unified atom of the protein model compounds at 23-25°C. Unfavorable interactions have positive α-values.

From these comparisons we deduce that the dramatically different effects of KCl vs. KGlu on SSB LLPS, especially at high salt where KGlu promotes but KCl inhibits LLPS, result from the very different non-Coulombic interactions of these salts with amide groups in the IDL regions of the C-terminal tails which are buried in amide-amide interactions that help drive SSB LLPS. KGlu interacts unfavorably with these amides in solution and therefore promotes amide-amide interactions in LLPS and NNN-cooperativity. By contrast, KCl interacts favorably with these amide groups in solution and therefore inhibits LLPS andNNN-cooperativity.

## Discussion

The formation of membraneless biomolecular condensates (phase separation) is a well-recognized, biologically important process (Banani et al., 2017, Alberti et al., 2019, Shin and Brangwynne, 2017, Holehouse and Pappu, 2018). Phase separation has been studied most widely and thought to be most relevant in eukaryotic systems. This is due to the fact that intrinsically disordered regions (IDRs) of proteins, that are effective drivers of phase separation, make up ∼40% of eukaryotic proteomes, but less than ∼5% of bacterial proteomes (van der Lee et al., 2014). However, it has been argued that membraneless biomolecular condensates are also important in bacteria (Cohan and Pappu, 2020).

It has been shown that the *E. coli* SSB tetramer can undergo phase separation at conditions that mimic physiological (20mM NaCl, 50 mM NaGlu, 5 mM Mg Acetate, pH 7.5) (Harami et al., 2020). Most experiments in that study were performed by varying SSB concentration at or near 25°C. In our experiments we monitored the turbidity of SSB solutions upon lowering the temperature and determined the temperature at which turbidity is first observed (T_PS_) (Amiram et al., 2011, Lyons et al., 2013, Cinar et al., 2019) (Fig.2). This allowed us to construct phase diagrams (coexistence curves) of T_PS_ vs [SSB] (Rubinstein, 2003, Posey et al., 2018, Falahati and Haji-Akbari, 2019, Alberti et al., 2019) to determine how they are affected by solution conditions (salt concentration and type).

The temperature-induced onset of turbidity is reversible in a narrow temperature range indicating that the cloud points are reasonable proxies for the actual T_bimodal_ below which the system separates into two coexisting phases (Thomson et al., 1987, Petsev et al., 2003, Rubinstein, 2003, Posey et al., 2018, Falahati and Haji-Akbari, 2019). We also note that this approach (starting with highly soluble SSB in the buffer alone) allows one to interrogate conditions of any mixture of salts at any concentration.

### *Ec*SSB IDL is required for phase separation

Whereas the roles of the tetrameric SSB core (DNA binding domain) and the conserved acidic tips of the C-termini (binding site for numerous metabolic proteins) have been well recognized, the role of the SSB linker (IDL) that connects the two regions is less clear. Our recent studies revealed that the IDL affects the relative stabilities of the two major binding modes as well as cooperative binding to ssDNA (Kozlov et al., 2015, Kozlov et al., 2017, Kozlov et al., 2019). As shown previously (Harami et al., 2020) and in this report, the IDL is also needed for phase separation. However, the IDL does not play a role in the interaction of SSB with at least four of its SIP partners (Shinn et al., 2019). In general bacterial SSBs possess a broad range of IDLs differing in length from 25 to 125 residues and amino acid composition (Kozlov et al., 2015, Cohan et al., 2021). A survey of 134 bacterial SSB proteins showed that independent of the length a majority of the IDL’s have low complexity compositions rich in glycines and are predicted to form globules (Kozlov et al., 2015). A statistically significant clustering of Gly residues was found to be conserved across three out of six SSB classes: actinobacteria, α-proteobacteria, and g-proteobacteria (Cohan et al., 2021). Recent bioinformatics analysis of more than 700 SSBs from 15 major phylogenetic groups of bacteria also showed that ∼70% have prion like regions (Harami et al., 2020). Harami et al. (Harami et al., 2020) also showed the importance of the C-terminal tail of *Ec*SSB for phase separation and found that removal of the conserved acidic tip diminishes but does not eliminate phase separation highlighting the importance of the IDL.

Here we demonstrate that reducing the number of SSB tails from four to two reduces its propensity to undergo phase separation. We also show that removal of the *Ec*SSB IDL to form SSB-ΔL eliminates phase separation in buffer containing 0.10 M KGlu. This highlights the importance of multi-valency in protein systems that undergo phase separation. Phase separation is also eliminated if the *E. coli* IDL is replaced with the *P. falciparum* IDL to form SSB-*EcPfEc*. *Pf*SSB itself also does not undergo phase separation under the same conditions. The *Ec*SSB and *Pf*SSB IDLs differ in length, charge and amino acid composition (Fig. 1D). The *Ec*-IDL is rich in Glycine (∼30%), which occur in a series of short segregated linear clusters, XGGX and XGGGX, where X represents Q, R, A,I and W (Fig. 1D). It has been shown that the presence of GG repeats within disordered low complexity regions, and when flanked by Arginine (RGG/RG), promote phase separation and/or self-assembly of many proteins (Chong et al., 2018, Murthy et al., 2021, Kar et al., 2021). Similarly, G flanked by Q or aliphatic/aromatic residues were also found to be important in driving phase separation (Chong et al., 2018, Murthy et al., 2021). This is the case for the *Ec*-IDL. In contrast, the *Pf*-IDL contains only 2 Gly in the entire IDL (Fig. 1D) and has 24 charged residues (17 anionic (D, E) and 7 cationic (K,R) charged residues in *Pf*-IDL vs 3 (1 anionic (E) and 2 cationic (R)) in *Ec*-IDL. We hypothesize that the *Pf*-IDL exhibits a lower propensity for phase separation, in particular because of the large number of carboxylates and the small number of glycines in its amino acid composition.

Previous atomistic simulations predict a compact globule conformation for the *Ec*SSB tails, whereas the *Pf*SSB tails are predicted to be more extended. This was supported by hydrodynamic parameters determined from sedimentation velocity experiments at low [NaCl] (Kozlov et al., 2015). These properties suggest that the *Ec*SSB tail should be more prone to intermolecular interactions promoting condensation. Thus, both the amino acid sequence, composition and the IDL charge influence the different phase behaviors of the *Ec* and *Pf* IDLs.

Interestingly, we found that some partial deletions within the *Ec*SSB IDL (SSB*Δ*151- 166; SSB*Δ*130-166) enhance the propensity of SSB to undergo phase separation, whereas a larger deletion (SSB*Δ*120-166) has the opposite effect and a total deletion of the IDL (SSB- ΔL) eliminates phase separation. These differences appear to be due to changes in the Glycine content of these partial deletions. First we note that for these partial deletions the percentage of Gly residues in the remaining IDL increases from 42% (SSB*Δ*151-166), to 53% (SSB*Δ*130-166) and to 55% (SSB*Δ*120-166) (Figure 1D) and the ratio of longer GGG vs shorter GG repeats also increases (2 vs 5, 2 vs 2, 1 vs 1, respectively). Gly-rich regions can drive self-assembly, especially if the Gly-rich regions or poly-Gly tracts are increased (Kar et al., 2021). This is consistent with our finding that the SSB*Δ*130-166 construct shows the highest tendency for phase separation (53% Gly content and 2 GG and 2 GGG repeats). We speculate that this construct has an optimal balance of overall Gly content and number of GG and GGG repeats that promote phase separation. However, the specificity of the residues flanking these repeats may also be important and the precise balance among these factors remains to be determined. Interestingly, atomistic simulations predict that all C terminal deletion peptides (Fig.1D) should adopt compact conformations similar to the full length *Ec*-tail (Kozlov et al., 2015), and, therefore should show some propensity to undergo phase separation.

Although we find that a peptide with the *Ec*-IDL sequence does not undergo phase separation even in the presence of high concentrations of KGlu, this might be due to an inability to obtain sufficiently high peptide concentrations, which never exceeded ∼800 µM. A simple calculation assuming a random conformation of Ec-tail suggests that the concentration of the four C-terminal tails in the vicinity of the tetrameric core could be as much as few millimolar (or higher, if more compact) (Kozlov et al., 2015). If these high concentrations are required to achieve phase separation, then this might explain the decrease in T_PS_ that we observe for the two tailed *Ec*SSB-LD-*Drl* (Fig. 4A). However, the need for multi-valency, achieved by the tetrameric nature of *Ec*SSB, is also likely to be an important contributor to LLPS.

In addition to the IDL interactions, we note that interactions between the conserved C-terminal acidic tip (*MDFDDDIPF*) and the tetrameric DNA binding core might also contribute to phase separation. In fact, all of the deletion constructs that undergo phase separation still contain the acidic tip which can weakly interact with the positively charged tetrameric core and therefore compete for ssDNA binding (Kozlov et al., 2010a, Su et al., 2014, Kozlov et al., 2015, Kozlov et al., 2017, Harami et al., 2020). These additional core- tip interactions appear to enhance SSB phase separation. Importantly, we find that an SSB construct in which the negative charges in the tip are reversed to positive charges, *MKFKKKIPF* (all four D replaced with K) does not undergo phase separation, suggesting that the tip-core interactions are important, although we cannot exclude that the charge reversal may influence linker-linker interactions.

### Effects of KGlu and KCl on Phase Separation Indicate the Significance of Amide-Amide Interactions but not Charge-Charge Interactions Involving *Ec*SSB Tails

Effects of salt concentration on phase separation have been examined for a number of systems, primarily using chloride salts (NaCl/KCl ) (Boyko et al., 2019, Boeynaems et al., 2017, Nott et al., 2015, Brady et al., 2017, Burke et al., 2015). In most cases, increases in [NaCl] or [KCl] (up to 0.3 M) decreased or even eliminated phase separation. However, the opposite effect was shown for FUS LC (Burke et al., 2015) and lysozyme (Taratuta et al., 1990). Only a limited number of studies have explored phase separation at higher salt concentrations (>0.5 M) (Krainer et al., 2021, Taratuta et al., 1990) or by varying the anion type (Taratuta et al., 1990, Mason et al., 2010, Boeynaems et al., 2017, Zhang and Cremer, 2009). These latter studies show that the driving forces for phase separation are reduced for anions that interact more favorably with protein groups (i.e., are on the “salting in” end of the Hofmeister series (Hofmeister, 1888, von Hippel and Schleich, 1969, Pegram and Record, 2007): F^-^> Cl^-^> Br^-^> ClO_4_^-^ >I^-^ >SCN^-^ (Taratuta et al., 1990, Zhang and Cremer, 2009, Mason et al., 2010) and HPO_4_ >CO_3_ >Cl (Boeynaems et al., 2017)). Similar effects of Hofmeister cations following the series Na^+^∼K^+^>Rb^+^>Cs^+^>Li^+^∼Ca^2+^ were shown for FUS protein (Krainer et al., 2021). Tsang et al. (Tsang et al., 2019) examined the effects of several salt types on phase separation of the Fragile X FMRP protein bound to RNA, but all were chloride salts.

Since Glu^-^ is the major monovalent cytoplasmic anion in *E. coli*, ranging in concentration from 0.03 to 0.25 molal (Richey et al., 1987, Record et al., 1998), we chose to compare the effects of KGlu vs KCl on *Ec*SSB phase separation. Replacement of chloride with glutamate enhances the binding affinities of many proteins for nucleic acids (Leirmo et al., 1987, Kontur et al., 2010, Deredge et al., 2010) including *Ec*SSB (Overman et al., 1988) and also affects SSB NNN cooperativity (Kozlov et al., 2017). Harami et al. (Harami et al., 2020) showed that an increase in [NaCl] eliminates phase separation, although phase separation is still observed at increased [NaGlu]. Here we examined the profiles of the low concentration arms of coexistence curves as influenced by [KGlu] and [KCl]. This approach provides a 3D phase diagram of T_PS_ as a function of two variables [SSB] and [salt].

Effects of Glu^-^ and Cl^-^ salts on T_PS_ result from differences in interaction of the ions of these salts with SSB in the condensate and in the dilute solution phase. In general, two types of salt ion-protein interactions contribute to these salt effects. Long-range Coulombic interactions of salt ions with protein charges are dominant at low salt concentration. These Coulombic interactions are primarily determined by salt ion valence, and therefore should be similar for the Cl^-^ and Glu^-^ salts studied here. Weak, short-range ion-specific interactions, like those of salt ions with hydrocarbon groups of proteins which are responsible for the Hofmeister ion series for protein processes, contribute more at high salt concentration where Coulombic effects are minimized.

For this discussion, we assume that salt effects on the transfer of an SSB tetramer from dilute solution to condensate arise from SSB-SSB interactions in the condensate that change the number of unneutralized SSB charges and/or the ASA and hydration of the SSB tetramer. Salt effects could also arise from the transfer of SSB itself, since the environment of the condensate is different from that of a dilute solution.

We hypothesize that the opposite effects of [KGlu] and [KCl] on *Ec*SSB phase separation result from the opposite directions of the noncoulombic, Hofmeister-like preferential interactions of these salts with amide groups of the IDL, especially with G backbone amides (17 per IDL) and perhaps also Q and N side chain amides (14 per IDL). KGlu favors both SSB LLPS (Fig 3) and SSB-DNA collapse (Fig. 5) (Kozlov et al., 2017) at all [KGlu] investigated (≥ 0.01 M), while KCl disfavors both these processes except at the lowest [KCl] investigated (0.01 – 0.03 M). This indicates that coulombic effects of these salts are not the major determinant of these salt dependences, and therefore that SSB-SSB charge-charge interactions are not a major determinant of LLPS or NNN cooperativity thermodynamics.

The insensitivity of T_PS_ to [KCl] at or below 0.03 M, a salt range where T_PS_ increases with increasing [KGlu], indicates compensation between a nonspecific coulombic effect of KCl on SSB- charge-charge interactions, favoring LLPS and increasing T_PS_, and an ion-specific effect of KCl, reducing T_PS_. Therefore the charge-charge interactions in LLPS appear to be between small numbers of like charges (e.g., the carboxylates at the tip of the SSB tails) and the coulombic effect of both salts screens these charges and reduces their unfavorable interactions. For KGlu, the predicted nonspecific Coulombic and ion-specific contributions to the observed increase in T_PS_ with increasing [KGlu] are not separable without a quantitative analysis (in preparation).

The large differences in effects of KGlu and KCl on T_PS_ are non-coulombic and anion-specific and result from differences in interactions of these salts with regions of SSB that are involved in LLPS and NNN cooperativity. The existence of these Hofmeister-like salt-specific differences means that SSB-SSB interactions in the high SSB concentration environment of the condensate reduce the water-accessible surface area (ASA) and/or hydration of SSB, resulting in release of water from this previously hydrated SSB surface. For KGlu to favor LLPS its ions must interact unfavorably in the solution phase with the SSB ASA that is involved in SSB-SSB interactions in the condensate. Conversely, for KCl to disfavor LLPS its ions must interact favorably in the solution phase with the SSB ASA that is involved in SSB-SSB interactions in the condensate.

From the *α*-values that quantify strengths of interaction of KGlu and KCl with a unit area of the different types of unified C, N and O atoms of SSB in Fig. 6, it is straightforward to deduce which SSB functional groups are most responsible for the opposite effects of [KGlu] and [KCl] on LLPS. Because SSB charge-charge interactions are of like charges and destabilizing for LLPS, no dehydration of these charged groups is expected in the SSB condensate. For this reason we focus on hydrocarbon and amide groups in the IDL of the *Ec*SSB tails to explain the different effects of KGlu vs. KCl.

(1) From Fig. 6 and Table S2, both salts interact unfavorably with aliphatic sp^3^C and aromatic sp^2^C, so burial of hydrocarbon ASA in the SSB condensate would be favored by addition of either salt and therefore would not explain the opposite effects of KGlu vs. KCl on LLPS.
(2) KGlu interacts unfavorably with amide sp^2^O and favorably with amide sp^2^N, while KCl interacts favorably with amide sp^2^O and unfavorably with amide sp^2^N. The most accessible amide groups on the SSB tails (those with the largest ASA) are the backbone amides of the 17 G residues and the 14 Q and N side chains. More than half of the amide ASA of G residues and about half the ASA of side chain amides is amide sp^2^O, and *α*-values (Fig. 6; Table S2) for the interactions of KGlu and KCl with amide sp^2^O are substantially larger in magnitude than *α*-values for interaction of these salts with amide sp^2^N. Therefore, the interactions with amide sp^2^O are predicted to dominate the salt-amide interaction. Because LLPS is more favorable in KGlu for the IDL deletion that removes residues 130-166 than for the full length IDL, and this Δ(130-166) variant has 10 G but only 1 Q and 1 N in 19 residues, it appears that the opposite interactions of the salts with amide groups of G residues are the most significant contributors to the salt effects, and therefore that burial and/or dehydration of G residues is a significant driving force for *Ec*SSB LLPS and, by extension, for NNN cooperativity.
(3) *Pf*SSB, as well as the chimeric SSB variant, *EcPfEc*SSB, do not undergo LLPS or display NNN cooperativity, even in KGlu. However, the *Pf*SSB tail also has a large number of N and Q amide side chains (29) but only two G residues, so the total number of amides is the same as in *Ec*SSB but with a very different distribution of G, N and Q. This also indicates that the most significant amide interactions in the *Ec*SSB condensate involve the G amide backbone. Furthermore, the *Pf*SSB tail also contains 21 anionic side chains vs 5 for *Ec*SSB, with only 7 *Pf*SSB cationic side chains. Hence, there would be a large coulombic cost to any tail-tail interactions in *Pf*SSB and *EcPfEc*SSB.

We propose that the large opposing effects of KGlu and KCl on LLPS provide a thermodynamic signature of extensive burial of amide residues in amide-amide interactions involving the many G residues of the *Ec*SSB tails. The fluidity of the condensate indicates that these amide interactions are very different than those involved in alpha helix or beta sheet formation where dehydration and direct hydrogen bonding interactions are involved. We therefore propose that these fluid amide-amide interactions are water-mediated, involving partial but not complete dehydration of amide sp^2^O and amide sp^2^N atoms.

Effects of amino acid composition and sequence on the ability of proteins and protein-nucleic acid complexes to undergo phase separation have been determined in numerous studies (Alberti et al., 2019, Posey et al., 2018, Choi et al., 2020, Martin et al., 2020). Here we show the dramatically different effects of the physiological salt KGlu and the laboratory salt KCl on SSB phase separation. We discover from the interactions of these salts with protein model compounds that the differences in their effects on SSB LLPS stems from differences in their interactions with amide groups, and consequently deduce that amide-amide interactions involving the many G residues and/or Q and N side chains of the IDL play a key role in SSB LLPS. Salt concentration and the nature of the salt ions clearly regulate LLPS in this system. This is a consideration in vivo, because the cytoplasmic [KGlu] varies over a wide range with growth osmolality (Richey et al., 1987, Cayley et al., 1991). Similar effects on phase separation of different salt types at high concentrations have been reported recently (Fetahaj et al., 2021). These salt effects are not ionic strength effects as in classical Debye-Huckel analyses because solutions of KCl and KGlu at the same concentration have the same ionic strength but have drastically different effects on SSB LLPS.

### SSB phase separation and NNN cooperative ssDNA binding are driven by the same forces

The *Ec*-IDL plays an important role in SSB binding to ssDNA, affecting the relative stabilities of the major binding modes and DNA binding cooperativity (Kozlov et al., 2015, Kozlov et al., 2017, Kozlov et al., 2019). Deletion of the IDL promotes the (SSB)_35_ binding mode and minimizes the effects of anion type (Kozlov et al., 2017). However, removal of the IDL or replacement of NaCl with NaGlu does not affect nearest-neighbor (NN) cooperativity in the (SSB)_65_ binding mode (Kozlov et al., 2019, Overman et al., 1988, Bujalowski and Lohman, 1987). At the same time, removal of the linker practically eliminates NN cooperativity in the (SSB)_35_ binding mode in low salt conditions (Ferrari et al., 1994, Kozlov et al., 2019).

The situation is strikingly different for the binding of SSB to polymeric ssDNA. In this case non-nearest-neighbor (NNN) cooperative interactions can occur among SSB tetramers distantly bound to the ssDNA, resulting in compaction/condensation of single nucleoprotein complexes (Bell et al., 2015, Kozlov et al., 2015, Kozlov et al., 2017). This was first observed using single molecule force microscopy for λ-ssDNA which undergoes additional compaction (beyond the expected compaction resulting from wrapping in the (SSB)_65_ mode) upon SSB binding at high sodium acetate concentrations (Bell et al., 2015). In fact, acetate is close to glutamate in the Hofmeister series and behaves similarly to glutamate. Indeed, we show here that the additional compaction of SSB-polymeric ssDNA complexes in KGlu is observed in ssDNA curtain experiments and is reversed when KGlu is replaced with KCl (0.30 M). Importantly, little additional compaction is detected for SSB-ΔL coated ssDNA indicating that this additional compaction is due to the interaction of KGlu with the IDL.

The cooperative formation of high binding density complexes was detected for the binding of SSB to M13 ssDNA using electrophoretic mobility shift assays at low [NaCl] (Lohman et al., 1986). It was manifested at low [NaCl] when less than saturating amounts of SSB are bound to the ssDNA in the form of bimodal distributions of the nucleoprotein complexes, with the faster moving band ascribed to the highly cooperative, fully saturated (compact) complex and the slower band reflecting the lower cooperativity (SSB)_65_ complexes. An increase in [NaCl] to 0.2-0.3 M eliminated these highly cooperative complexes. These observations are supported by sedimentation velocity experiments (Kozlov et al., 2015, Kozlov et al., 2017) (Fig.S3), which demonstrate: (1) at low [salt] (10 mM) bimodal distributions (NNN cooperativity) are not affected by the type of salt; (2) when [salt] increases to 0.20 M NaCl/KCl, the bimodal distributions are eliminated, but remain in the presence of even 0.50 M KGlu; (3) removal/replacement of the *Ec*SSB IDL totally eliminates high cooperativity in all salt conditions; (4) deletions within the IDL and reduction to two tails (SSB-LD-*Drl*) do not eliminate bimodal distributions.

These comparisons indicate that the same conditions that favor NNN cooperative binding of SSB on ssDNA also favor phase separation of SSB in the absence of DNA (Fig. 7), suggesting that both processes are governed by the same forces and both driven by linker-mediated interactions (Kozlov et al., 2010a, Su et al., 2014, Kozlov et al., 2015, Kozlov et al., 2017, Harami et al., 2020). Yet, we and Harami et al. (Harami et al., 2020) show that binding of even short lengths of ssDNA to SSB tetramers eliminates phase separation. However, when SSB is bound to polymeric ssDNA, phase separation can be replaced by NNN cooperative binding of SSB to ssDNA. In this sense, NNN cooperative binding of SSB to polymeric ssDNA and phase separation of SSB alone can be viewed as competitive processes. Since phase separation does not occur for SSB bound to ssDNA, the binding of SSB to a polymeric ssDNA causes collapse of a single DNA molecule.

**Fig. 7.**
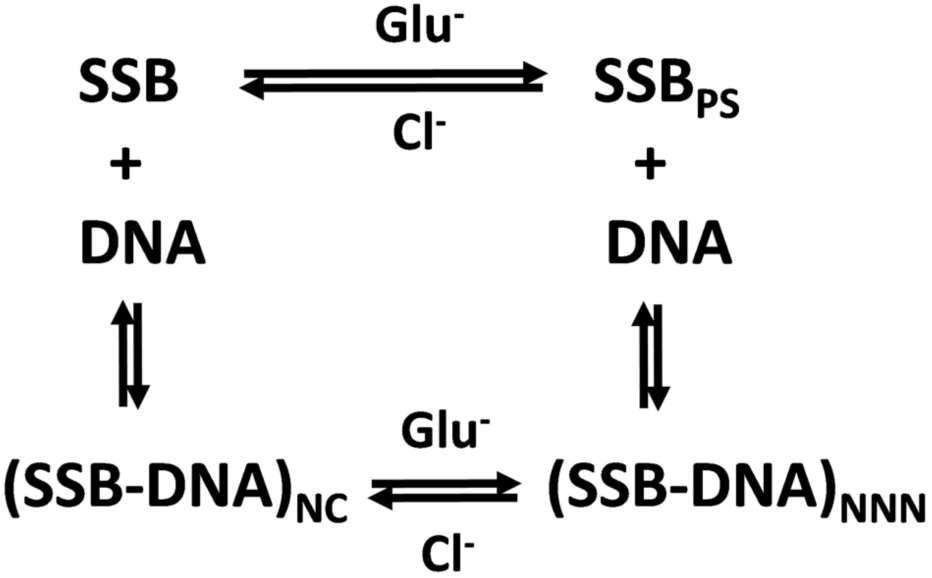
SSB LLPS and NNN cooperativity on polymeric ssDNA are driven by the same forces. This scheme shows the various states of SSB and its complexes with ssDNA: SSB is free SSB tetramer; SSB_PS_ is the SSB tetramer in the dense condensate; (SSB-DNA)_NC_ is SSB bound non-cooperatively to polymeric ssDNA; (SSB-DNA)_NNN_ is SSB bound with non-nearest neighbor cooperativity to polymeric ssDNA. Glutamate promotes and chloride inhibits both LLPS of free SSB as well as non-nearest-neighbor (NNN) cooperative complexes on polymeric ssDNA.

### Biological implications of SSB phase separation

It has been proposed that the propensity of *E. coli* SSB alone to undergo phase separation and the loss of this ability upon binding ssDNA implicates phase separation of SSB *in vivo* as a mechanism to sequester SSB until it is needed for DNA metabolism (Harami et al., 2020, Yu et al., 2016). First, highly concentrated/condensed SSB can be easily delivered to ssDNA. SSB has been reported to localize near the cell membrane (Zhao et al., 2019), as well as at replication forks (Spenkelink et al., 2019). Second, it is possible that SSB condensates may serve as vehicles to deliver myriad SSB interacting proteins (SIPs) to their places of action on DNA. SIPs usually interact weakly (K_d_∼0.1- 1µM) with SSB through the conserved acidic tip, (Kozlov et al., 2010b, Zhou et al., 2011, Shinn et al., 2019). However, in SSB condensates SIP accumulation should increase due to the high SSB concentrations (∼ 4 mM) (Harami et al., 2020). In fact, Harami et al. (Harami et al., 2020) showed that RecQ is enriched in SSB condensates *in vivo*. Third, dissolution of the SSB condensates upon interaction with DNA could result in the immediate coating of the ssDNA with SSB and result in the release of the SIPs to function on ssDNA. It is important to note that the loss of the ability of SSB bound to ssDNA to undergo phase separation is replaced by the ability of a single polymeric ssDNA molecule, when saturated with SSB, to undergo compaction/condensation via NNN cooperative interactions of DNA bound SSB tetramers. This, in turn, could bring together different proteins bound exclusively to SSB or ssDNA for further action.

## Materials and Methods

### Reagents and buffers

Buffers were prepared with reagent grade chemicals and distilled water treated with a Milli Q (Millipore, Bedford, MA) water purification system. Buffer T is 10 mM Tris, pH 8.1 (25°C), 0.1 mM Na_3_EDTA, Buffer P is 10 mM phosphate, pH 7.5 (25°C), 0.1 mM Na_3_EDTA. The final concentrations of monovalent salts (KGlu, KCl and NaCl, Sigma-Aldrich, reagent grade) in the solutions were achieved by mixing with solutions of Buffer T or Buffer P containing 1 M KGlu, 2 M KCl or 2 M NaCl. Single molecule imaging buffers (I) and (II) are 10 mM Tris-HCl, pH 7.5, 1 mM DTT, and 0.2 mg/ml BSA (NEB B9000S) and 10 mM Tris-HCl, pH 8.1, 0.5 mM 2-mercaptoethanol, respectively.

#### Plasmodium falciparum

SSB *(Pf*SSB), *E. coli* SSB protein (*Ec*SSB) and its tail variants (see Fig. 1) were expressed and purified as described (Antony et al., 2012, Kozlov et al., 2015) including a new variant, SSB-Ktip in which 4 Asp in the tip sequence were replaced with 4 Lys. All SSB proteins in this study form stable tetramers under all solution conditions used in this study as determined by sedimentation velocity (Antony et al., 2012, Kozlov et al., 2015). SSB-LD-*Drl* is an SSB dimer construct in which two OB folds of each monomer are covalently linked using a 23 amino acid linker from *Dr*SSB (Fig.1C(iii)) as described (Antony et al., 2013) and thus possesses only two C-terminal tails (Fig.1C(iii)). Protein concentrations were determined spectrophotometrically (Lohman and Overman, 1985) (buffer T, 0.20 M NaCl) using *ε*_280_=1.13 *×* 10^5^ M^-1^ cm^-1^ for wtSSB, SSB*Δ*151-166 and SSBKtip; *ε*_280_=8.98 *×* 10^4^ M^-1^ cm^-1^ for SSB*Δ*130-166, SSB*Δ*120-166, SSB-*ΔL* and SSB-*EcPfEc*; *ε*_280_=9.58 *×* 10^4^ M^-1^ cm^-1^ for *Pf*SSB; and *ε*_280_=1.01 *×* 10^5^ M^-1^ cm^-1^ for SSB-LD-*Drl* and are reported as SSB tetramer concentrations (or dimers in the case of SSB-LD-*Drl*).

Single stranded M13 mp18 DNA used for sedimentation velocity experiments was from New England Biolabs (Catalog #N4040S). The concentration was determined spectrophotometrically in buffer T + 0.10 M NaCl using *ε*_259_ = 7370 M^-1^ cm^-1^ (nucleotide)(Berkowitz and Day, 1974). Low-complexity single stranded DNA substrates used for the ssDNA curtains were prepared as described (Schaub et al., 2018, Zhang et al.,AAA AAA AGA AAA GAA GG) and 4.5 μM biotinylated primer oligo IF238 (5/Biosg/TC TCC TCC TTC T) were annealed in T4 ligase reaction buffer (NEB B0202S). The mixture was heated to 75°C for 5 min and cooled to 4°C at a rate of −1°C min^−1^. Annealed circles were ligated with the addition of 1 μL of T4 DNA ligase (NEB M0202S) at room temperature for ∼4 hours. Low-complexity ssDNA was synthesized in phi29 DNA polymerase reaction buffer (NEB M0269S), 500 μM dCTP and dTTP (NEB N0446S), 0.2 mg mL^-1^ BSA (NEB B9000S), 10 nM annealed circles, and 100 nM of home-made phi29 DNA polymerase. The solution was mixed and immediately injected into the flow cell and incubated at 30°C for ∼30 min. ssDNA synthesis was quenched by removing excess nucleotides and polymerase with imaging buffer (I). ssDNA was end-labeled with mouse anti-dsDNA primary antibody (Thermo MA1-35346) followed by Alexa488-labeled goat anti-mouse secondary antibody (Thermo A28175) in the flow cell.

### Turbidity measurements

Turbidity measurements were performed using a Cary-100 spectrophotometer (Agilent, Santa Clara, CA) with a Temperature-Controller 550 and a 8x6 muliti-cell block- 750 thermostat accessory. Typically, low volume (140 µL) cells (1 cm pathway) were filled with solutions containing protein and varied buffer/salt composition at a temperature above the apparent phase separation transition temperature, T_PS_, where the solution is totally transparent. Phase separation was monitored by following the increase in turbidity (light scattering) at 600 nm upon continuous decrease of the temperature (0.2 °C/min). The reversibility of the transitions was monitored by increasing the temperature at a rate of 0.2- 0.3°C/min.

### Analytical sedimentation

Sedimentation velocity experiments were performed as described (Kozlov et al., 2015, Kozlov et al., 2017, Kozlov et al., 2019) with an Optima XL-A analytical ultracentrifuge and An50Ti rotor (Beckman Instruments, Fullerton, CA) at 15000 rpm in buffer T, 25°C, with the salt concentration and type as indicated. A constant DNA concentration (typically 25 μM) was used and the protein/DNA ratios are indicated as R_65_=65x[P_tot_]/[DNA(nts)_tot_], where [P_tot_] is the total SSB tetramer concentration and [DNA(nts)_tot_] is the total DNA concentration in nucleotides. The absorbance was monitored at 260 nm, which mainly reflects the absorbance of the ssDNA, as the contribution of SSB to the absorbance at 260 nm is very small compared to the DNA at protein/DNA ratio R_65_ ∼0.6 used in this study (Kozlov et al., 2015, Kozlov et al., 2017, Kozlov et al., 2019). Data were analyzed using SEDFIT (www.analyticalultracentrifugation.com) to obtain c(s) distributions (Dam and Schuck, 2004). The densities and viscosities at 25°C were calculated using SEDNTERP for KCl/NaCl solutions and from van Holst et al. (van Holst et al., 2008) for KGlu solutions.

### Single-molecule Fluorescence Microscopy

Flow cells were prepared as described (Soniat et al., 2017, Schaub et al., 2018). Briefly, a 4-mm-wide, 100-μm-high flow channel was constructed between a glass coverslip (VWR 48393 059) and a custom-made flow cell containing 1−2-μm-wide chromium barriers using two-sided tape (3M 665). Single-molecule fluorescent images were collected with an inverted Nikon Ti-E microscope using prism based TIRF microscopy. The sample was illuminated with a 488 nm laser (Coherent Sapphire; 5 mW at front prism face). Fluorescent imaging was recorded using electron-multiplying charge-coupled device (EMCCD) camera (Andor iXon DU897). Subsequent images were exported as uncompressed TIFF stacks for further analysis.

DNA ends were tracked using custom written FIJI scripts. Briefly, the intensity of fluorescent spot located at the ssDNA end was fit to a two-dimensional Gaussian for sub-pixel particle localization. The trajectory of the center of the Gaussian was then plotted as a function of time. The plateaus in length change were then averaged together to reveal the corresponding length at different conditions. Kymographs were generated by taking a single-pixel wide section for each individual DNA molecule. 200 nM protein in imaging buffer (I) containing BSA was injected into the flow cell to generate *Ec*SSB-coated ssDNA or *Ec*SSB-ΔL-coated ssDNA. Buffer exchange experiments were conducted in either imaging buffer (I) or imaging buffer (I) supplemented with 300 mM KCl or 300 mM KGlu. All experiments were conducted at 1 mL min^-1^ flow rate at 37°C. Data were collected with a 50- ms exposure at a 5 second frame rate.

Fluorescence imaging experiments (Fig. S1) were conducted with an Olympus IX83 Cell^tirf^ equipped with three lasers, an image splitter (Olympus), an ANDOR iXon Ultra 897U EMCCD camera (Oxford Instruments), a temperature controlled slide holder (Warner Instruments TC-324C), and an objective heater (Warner Instruments TC-124A). A glass flow-channel was coated with PEG-5000 surface to prevent non-specific bindings. The experiments were performed in imaging buffer-(II) at 5 μM unlabeled SSB (+20 nM of SSB labeled with Cy5 at position A122C in the tail as described (Sokoloski et al., 2016)). The sample was excited with a 640-nm laser, and the emission was passed through an ET700/75M filter (Chroma). The data was recorded with a 30-ms exposure for 2000 frames using the Olympus cellSens Dimension software.

Confocal fluorescence measurements (Fig. S2) were performed on a Picoquant MT200 instrument (Picoquant, Germany) using an Olympus microscope (Olympus IX-73, Japan) equipped with water immersion objective (60x 1.2 UPlanSApo Superapochromat, Olympus, Japan). The Alexa 555 fluorescence was excited using a 485 nm pulsed laser (LDH PC- 485, Picoquant, Germany) and emitted photons were collected through the objective and separated according to polarization using a polarizer beam splitter cube (Ealing, Scotts Valley, CA, USA) and further refined by a 642 ± 40 nm bandpass filter (E642/80m, Chroma, Bellows Falls, VT, USA). All measurements were performed in uncoated polymer coverslip cuvettes (30 μl per well) (Ibidi, Germany) at 23 ± 1°C in a temperature-controlled room. Imaging was performed using both XY and Z monodirectional scanning with 1 ms collecting steps with 256x256 pixel resolution. The experiments were performed in imaging buffer-(II) at 4 μM unlabeled SSB (+20 nM of SSB labeled with Alexa Fluor 555 at position A122C in the tail as described (Sokoloski et al., 2016)).

### Vapor pressure osmometry

Osmolalities of solutions of protein model compounds and/or KCl were measured on Wescor Vapro 5600 Vapor Pressure Osmometers (VPO) calibrated as previously described (Cheng et al., 2016, Knowles et al., 2015, Cheng et al., 2017). Differences in osmolality (*ΔOsm*(*m*_2_,*m*_3_)) between a three-component solution (*Osm*(*m*_2_,*m*_3_)) and the corresponding two-component solutions (*Osm*(*m*_2_), *Osm*(*m*_3_)) were calculated using Eq. 1:

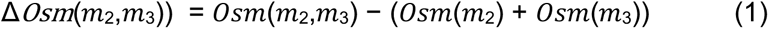

These Δ*Osm*(*m*_2_,*m*_3_) quantify the free energy consequences of interactions between the two solutes in water, as shown in Eq. 2:

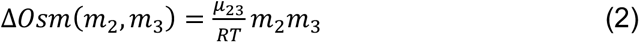

where μ_23_ is the chemical potential partial derivative (∂μ_2_/∂m_3_)_T.P,m2_ which quantifies the preferential interaction of solute 3 (salt) with solute 2 (model compound), relative to interactions with water (Robinson and Stokes, 1961, Capp et al., 2009, Cheng et al., 2020).

Hence the slope of a plot of Δ*Os*(*m*_2_,*m*_3_) vs. *m*_2_*m*_3_ is μ_23_/RT. In Eq. 2, the product *m*2*m*3 is the probability of an interaction of species 2 and 3, and 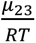 is the intrinsic strength of that interaction.

### Solubility

The molal-scale concentration of naphthalene in the <u>s</u>aturated <u>s</u>olution (*m*^77^) at 25 °C was determined as a function of KCl concentration by previously described methods (Knowles et al., 2015, Cheng et al., 2017). Values were normalized by the fitted solubility in the absence of KCl and the natural logarithm of the normalized solubility was plotted vs. KCl molality, *m_3_*. The limiting value of the salt-naphthalene preferential interaction coefficient μ_23_ (i.e. the solubility “*m*-value”) is determined from the initial slope of this plot:

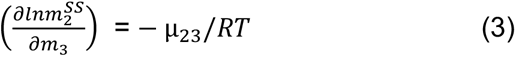

### Intrinsic Strengths of Interaction of KCl with Unified O, N, and C Atoms of Protein Groups (α-Values)

Analysis of the set of μ_23_ values for interactions of KCl with protein model compounds yields intrinsic strengths of interaction (designated α-values) of KCl with a unit area (ASA) of the various types (hybridization states, functional group context) of C, N, and O unified atoms of proteins. (Unified C, N and O atoms include all covalently bonded hydrogens). Details of this analysis for salts and other solutes have been described (Cheng et al., 2016, Cheng et al., 2017). A negative (positive) *α*-value indicates a favorable (unfavorable) interaction of KCl with that atom-type. This analysis has been used to quantify interactions of a wide range of biochemical solutes with unified atoms of proteins and nucleobases, and thereby predict or interpret effects of these solutes on protein and nucleic acid processes (Sengupta et al., 2016, Sengupta et al., 2017). α-Values are obtained by dissection of μ_23_ values into additive contributions *α_i_ASA_i_* where ASA_i_ is the water accessible surface area of one of the six types of unified atom (where the index i (1 ≤ i ≤ 6) represents aliphatic sp^3^C, aromatic and amide sp^2^C, amide sp^2^N, cationic sp^2^N and sp^3^N, amide sp^2^O, anionic (carboxylate) sp^2^O) on the model compounds.

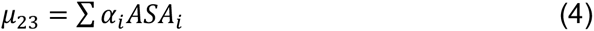

These ASA values have been reported (Cheng et al., 2016, Cheng et al., 2017). The set of μ_23_ values (19 in this case) is much larger than the number of *α_i_* values (6) being determined, testing the assumption of additivity in Eq. 4 and yielding unique determinations of the *α_i_* values by multiple linear regression. Values of *α_i_* for KGlu interactions with these six types of protein unified atoms have been reported (Cheng et al., 2016). Reported uncertainties for µ_23_ are one standard deviation based on the fitting error, and for α-values are the propagated uncertainty (Knowles et al., 2015).

## Acknowledgments

We thank R. Galletto for use of his Cary-100 spectrophotometer for turbidity measurements and J. Incicco and A. Soranno for help with imaging experiments. We thank Rohit Pappu for stimulating discussions, advice, enlightening comments and encouragement, and Kacey Mersch for discussions and comments on the ms. This research was supported in part by the NIH (R01 GM30498 and R35 GM136632 to TML) (R35 GM118100 to MTR), and NSF (CAREER award 1453358 to IJF).

## Abbreviations

LLPS: liquid-liquid phase separation
SSB: single stranded DNA binding protein
IDP: intrinsically disordered protein
IDL: intrinsically disordered linker
SIP: SSB interacting protein
ssDNA: single stranded DNA

## Author Contributions

AGK, TML, MTR and IJF designed the research; EAW made the SSB variants; MKS, BN performed imaging experiments; AGK performed and analyzed the experiments, XC performed and analyzed the osmometry experiments, IS and EZ analyzed the osmometry experiments, HZ performed and analyzed the single-molecule experiments. IJF supervised the single-molecule work. AGK, TML and MTR wrote the manuscript.

## Supplemental Information

**Figure S1.**
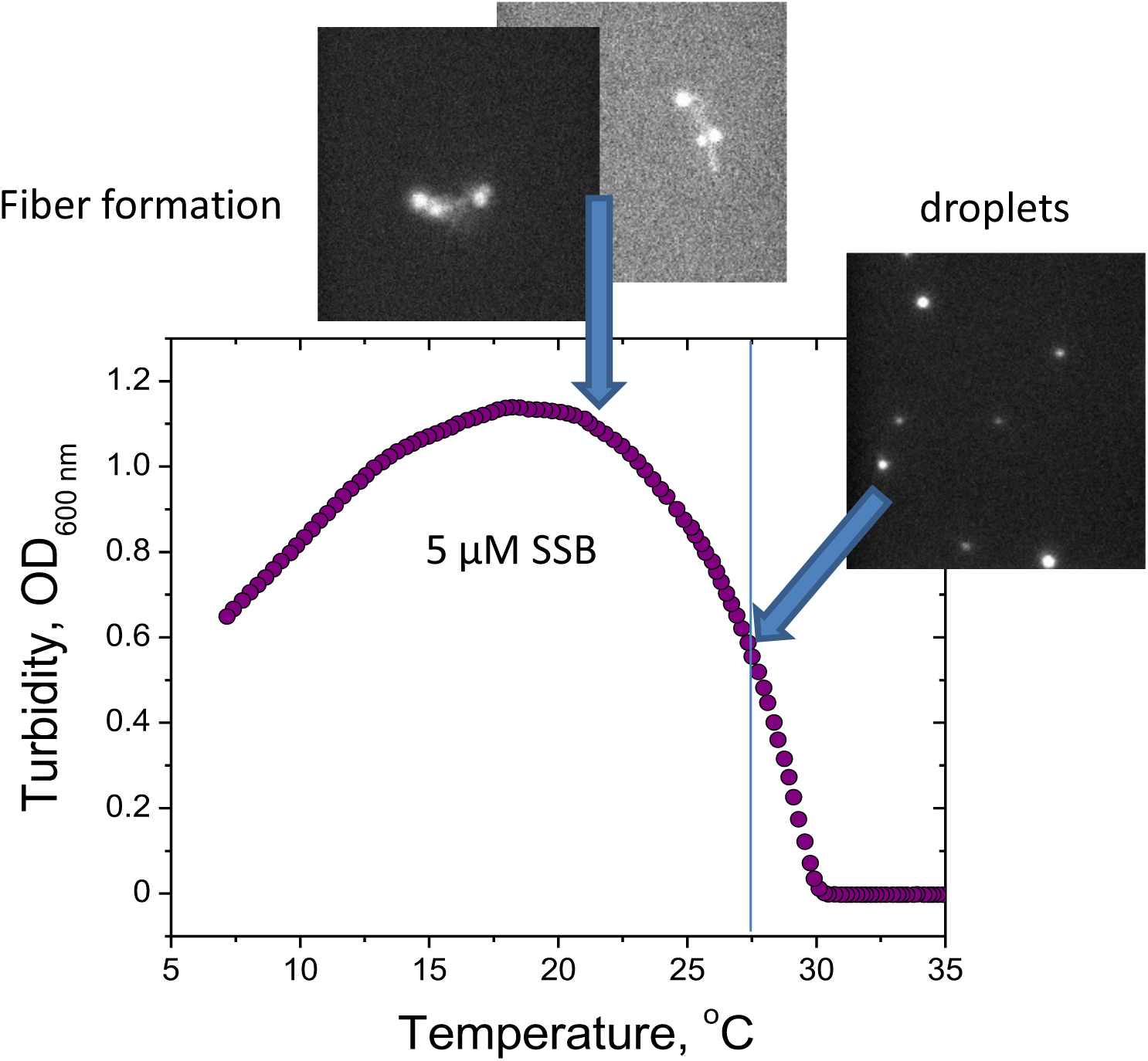
Microscopic observations support formation of liquid phase droplets in the range of linear increase of turbidity and formation of solid phase at the maximum of turbidity curve. Change in turbidity (OD at 600 nm) of a wtSSB protein solution (5 µM) (buffer T, 0.20 M KGlu) measured upon decreasing temperature from 35°C to 3°C and reflecting formation liquid-liquid phase at temperatures below T_PS_=30.1°C. Superimposed with the turbidity curve are microscopic images of a 5 µM SSB solution containing 20 nM of SSB labeled with Cy5 (buffer T, 0.20 M KGlu ) at 27.5°C and 22.5°C using TIRF confocal microscope (see Materials and Methods) and reflecting appearance of liquid droplets in the linear part of turbidity curve and fibril like structures at the maximum of turbidity change, respectively.

**Figure S2.**
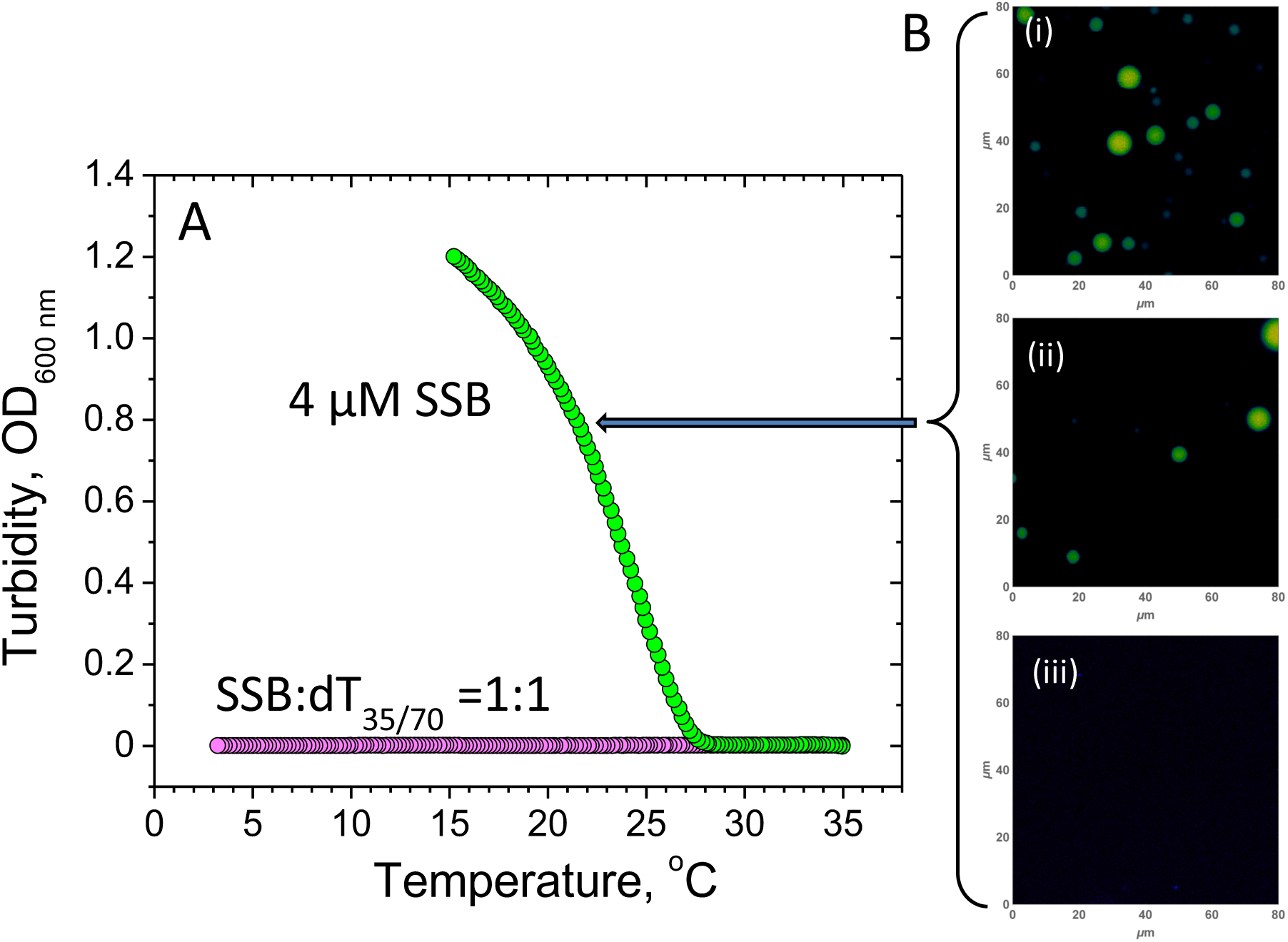
DNA binding to SSB inhibits phase separation. **(A)** Turbidity curves for SSB alone (4 µM, green) and for preformed SSB:dT_35/70_ =1:1 complexes (15 µM, magenta) obtained upon decreasing temperature in the range from 35°C to 3°C with a rate of 0.2 °C/min (buffer T, 0.1M KGlu). **(B)** Microscopic images of 4 µM solution of SSB (+ 20 nM of SSB labeled with Alexa 555; exc: 485 nm, em: 642±40 nm) obtained for (i)- SSB alone; (ii)- and upon addition of 2 µM dT_35_, and (iii)- addition of 4 µM dT_35_ (buffer T, 0.1M KGlu). The images were obtained at room temperature, ∼23_o_C, in the region of intermediate turbidity (as indicated by the arrow in panel A).

**Figure S3.**
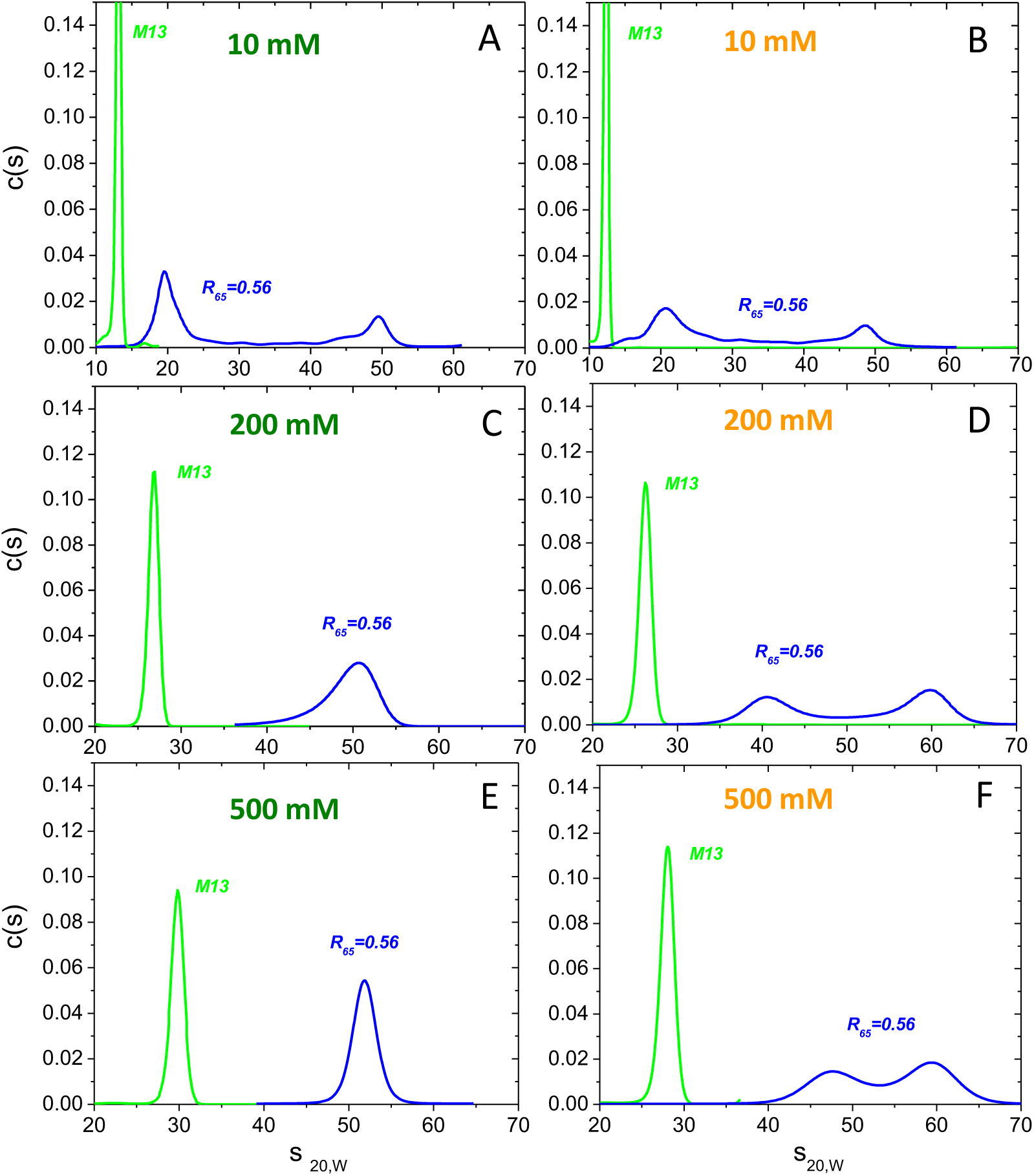
Salt concentration and type regulate non nearest-neighbor (NNN) cooperative binding of wtSSB to M13-ssDNA. (modified from Fig. 3 of Kozlov et al(Kozlov et al., 2017)). Highly cooperative SSB-ssDNA binding persists in high [KGlu], but is diminished in high [KCl]. Representative sedimentation velocity c(s) distributions converted to 20 °C, water conditions for wtSSB-M13ssDNA complexes at protein to DNA ratio: R_65_ = 0.56 (blue), where R_65_ = [SSB_tetr,tot_] × 65/[M13ssDNA_nts,tot_]. M13ssDNA alone (25 μM nts) is shown in green. (a) – 10 mM KCl, (b) – 10 mM KGlu, (c) – 0.20 M KCl, (d) – 0.20 M KGlu, (e) – 0.50 M KCl and (f) – 0.50 M KGlu.

**Figure S4.**
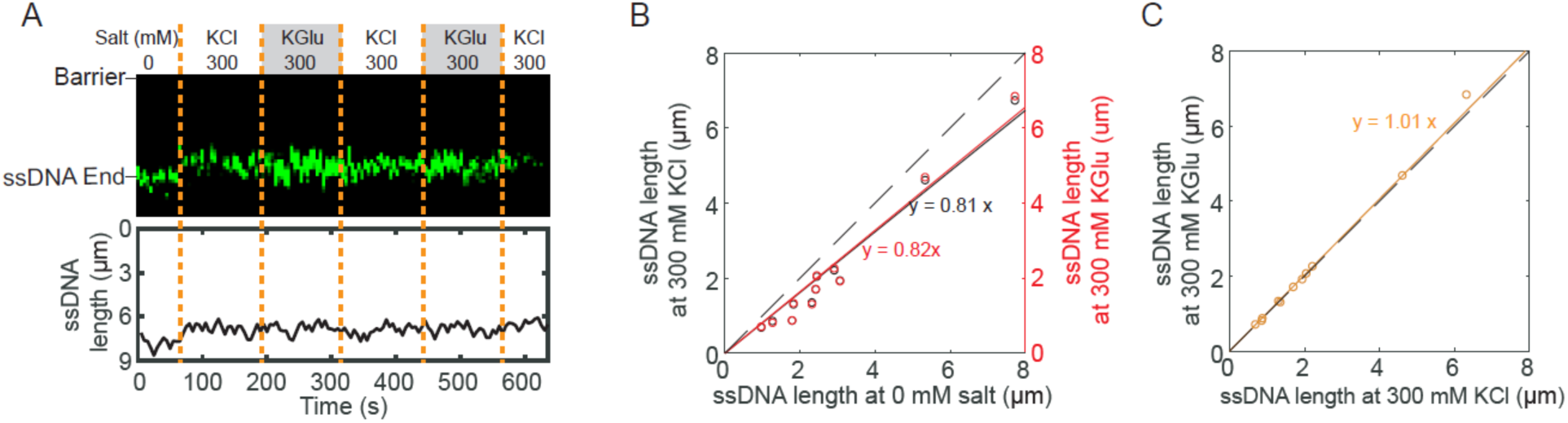
Change in salt concentration and type have little or no effect on the length of unbound ssDNA. (**A**) Representative kymograph (top) and single-particle tracking (bottom) showing the effects of salt on ssDNA end (green). Dashed orange lines denote when the buffer was switched. (**B**) Correlation between ssDNA lengths at 0 and 300 mM KCl (black), and at 0 and 300 mM KGlu (red). The solid lines are a linear fit to the data (N = 11 molecules). The dashed line represents a slope of 1. (**C**) Correlation between ssDNA lengths at 300 mM KCl and 300 mM KGlu (orange).

**Figure S5.**
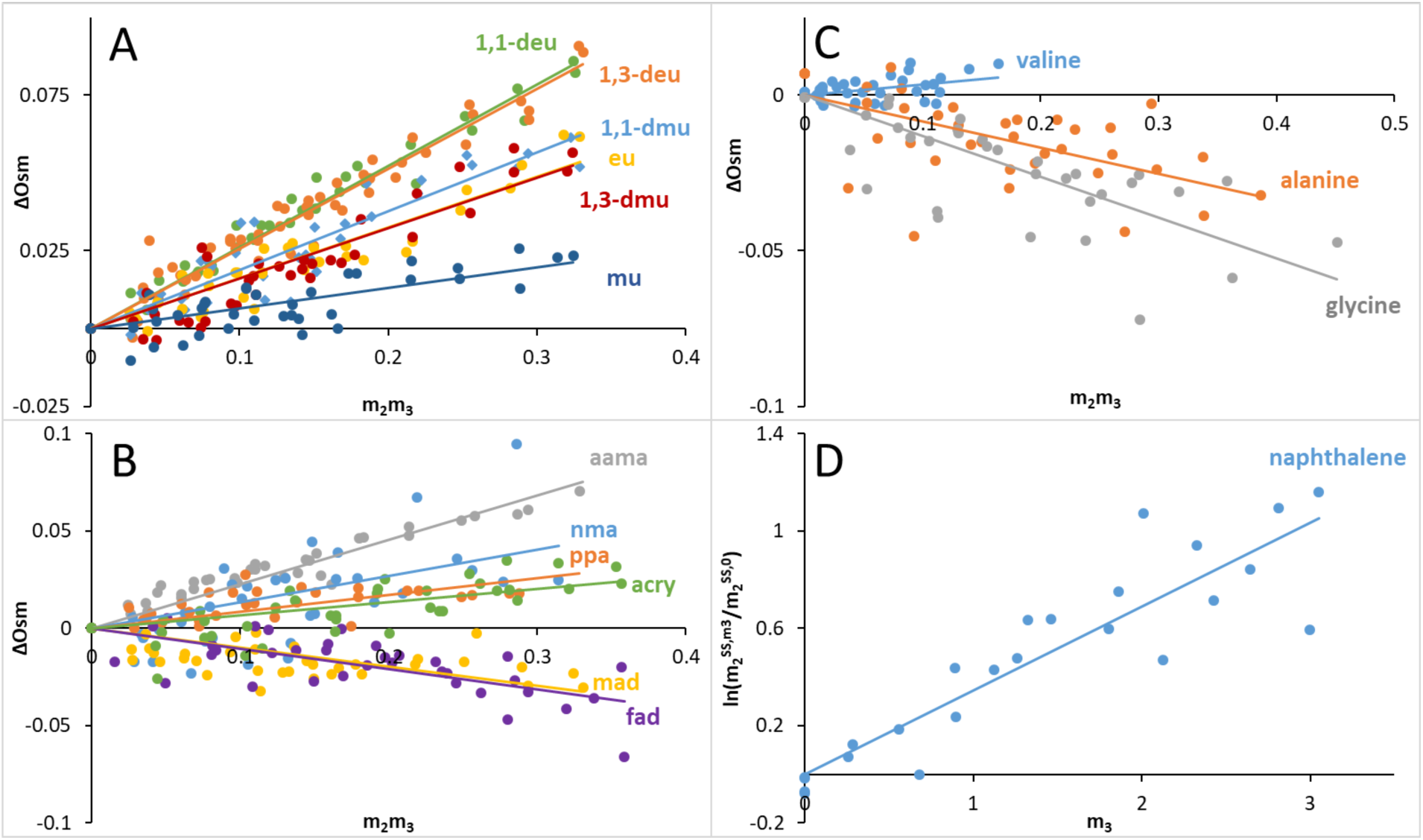
Determinations of μ_23_ for Interactions of KCl with Protein Model Compounds by Osmometry and Solubility Assays. In Panels A – C, osmolality differences ΔOsm (defined in Eq. 1) at 23 °C are plotted as a function of the product of molalities of KCl (m_3_) and the model compound (m_2_) according to Eq. 2 and μ_23_ is obtained from the slope. In Panel D, the logarithm of the normalized solubility of naphthalene (m2^SS^/m2^SS,0^) at 25 °C is plotted vs the molal concentration of KCl (m_3_) according to Eq 3 and μ_23_ is obtained from the slope.

**Figure S6.**
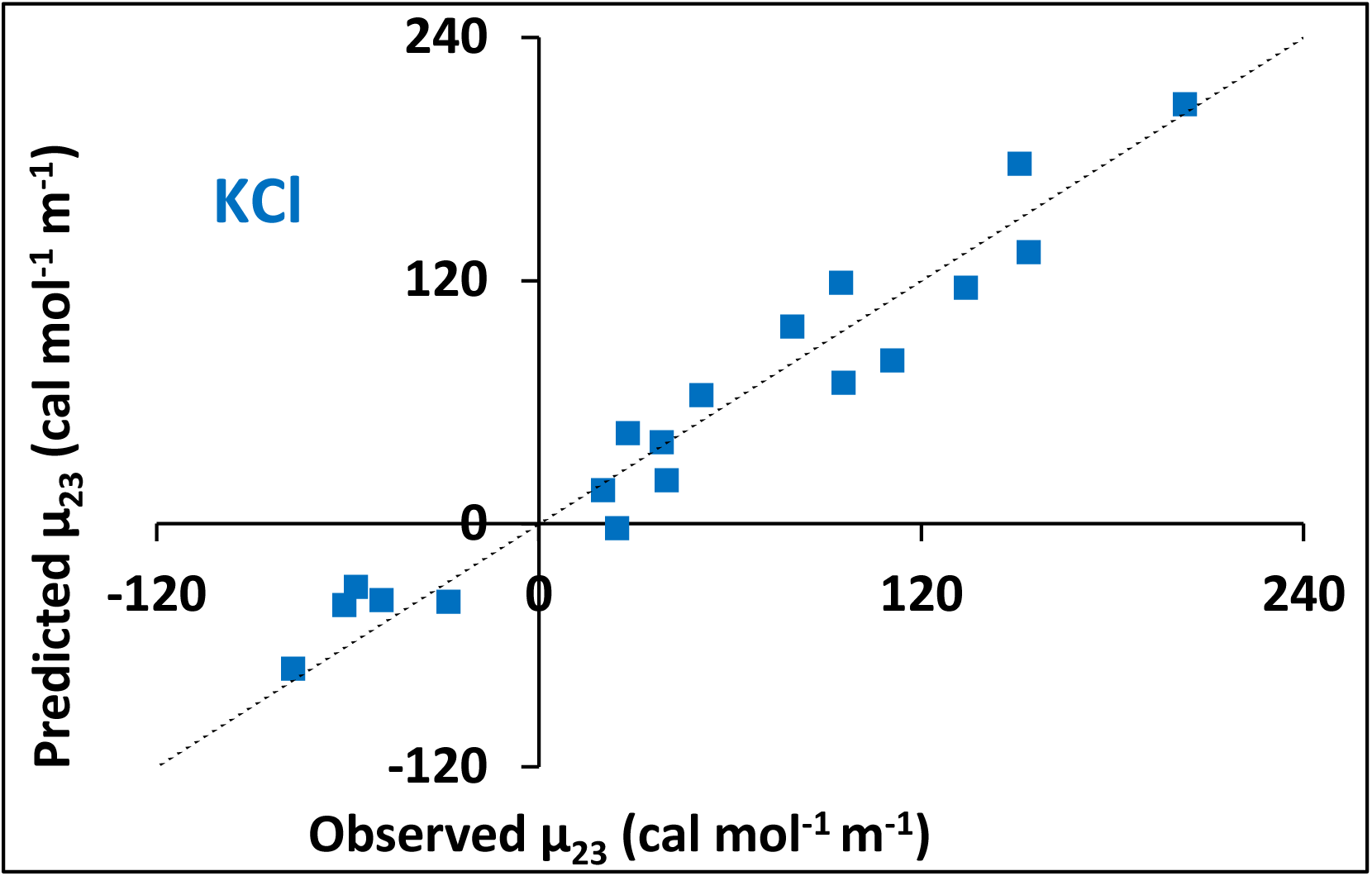
Predicted vs Observed μ_23_ Values for KCl. Values are listed in table S1. The dashed line represents agreement between predicted and observed μ_23_ values.

**Table S1.** Values of µ23 at 23 °C for KCl- model compound interactions and comparison with KGlu (Cheng et al., 2016). Three compounds with primarily amide ASA are listed first. Other compounds are listed in order from the most favorable to the most unfavorable interaction with KCl. Units of µ23 are cal mol^-1^ m^-1^.

| Model<br>Compound | KCl |  | KGlu |  |
| --- | --- | --- | --- | --- |
|  | Observed | Predicted | Observed | Predicted |
| formamide | -61 ± 5 | -40 ± 6 | * | * |
| malonamide | -57 ± 5 | -31 ± 10 | 48 ± 2 | 77 ± 13 |
| urea | -28 ± 3 | -39 ± 10 | 31 ± 2 | -6 ± 11 |
| glycine | -77 ± 6 | -72 ± 10 | -50 ± 4 | -38 ± 8 |
| alanine | -49 ± 6 | -38 ± 10 | 29 ± 2 | 24 ± 8 |
| valine | 20 ± 4 | 16 ± 11 | 117 ± 5 | 109 ± 8 |
| proline | 25 ± 4 | -3 ± 11 | 161 ± 4 | 160 ± 7 |
| glycine betaine | 28 ± 1 | 45 ± 11 | 292 ± 4 | 294 ± 7 |
| methylurea | 39 ± 3 | 40 ± 7 | 111 ± 6 | 121 ± 8 |
| acrylamide | 40 ± 5 | 21 ± 5 | * | * |
| propionamide | 51 ± 5 | 63 ± 7 | 174 ± 6 | 176 ± 7 |
| methylacetamide | 80 ± 12 | 97 ± 7 | * | * |
| 1,3-dmu | 95 ± 4 | 119 ± 8 | 249 ± 5 | 248 ± 6 |
| ethylurea | 96 ± 3 | 69 ± 8 | * | * |
| 1,1-dmu | 111 ± 4 | 80 ± 7 | * | * |
| aama | 134 ± 4 | 116 ± 11 | 406 ± 6 | 390 ± 11 |
| 1,3-deu | 151 ± 3 | 177 ± 10 | 348 ± 7 | 350 ± 7 |
| 1,1-deu | 154 ± 2 | 134 ± 9 | 284 ± 4 | 292 ± 8 |
| benzene | * | * | 291 ± 11 | 265 ± 8 |
| naphthalene | 203 ± 12 | 207 ± 4 | 322 ± 14 | 342 ± 10 |

\*not determined

**Table S2.**
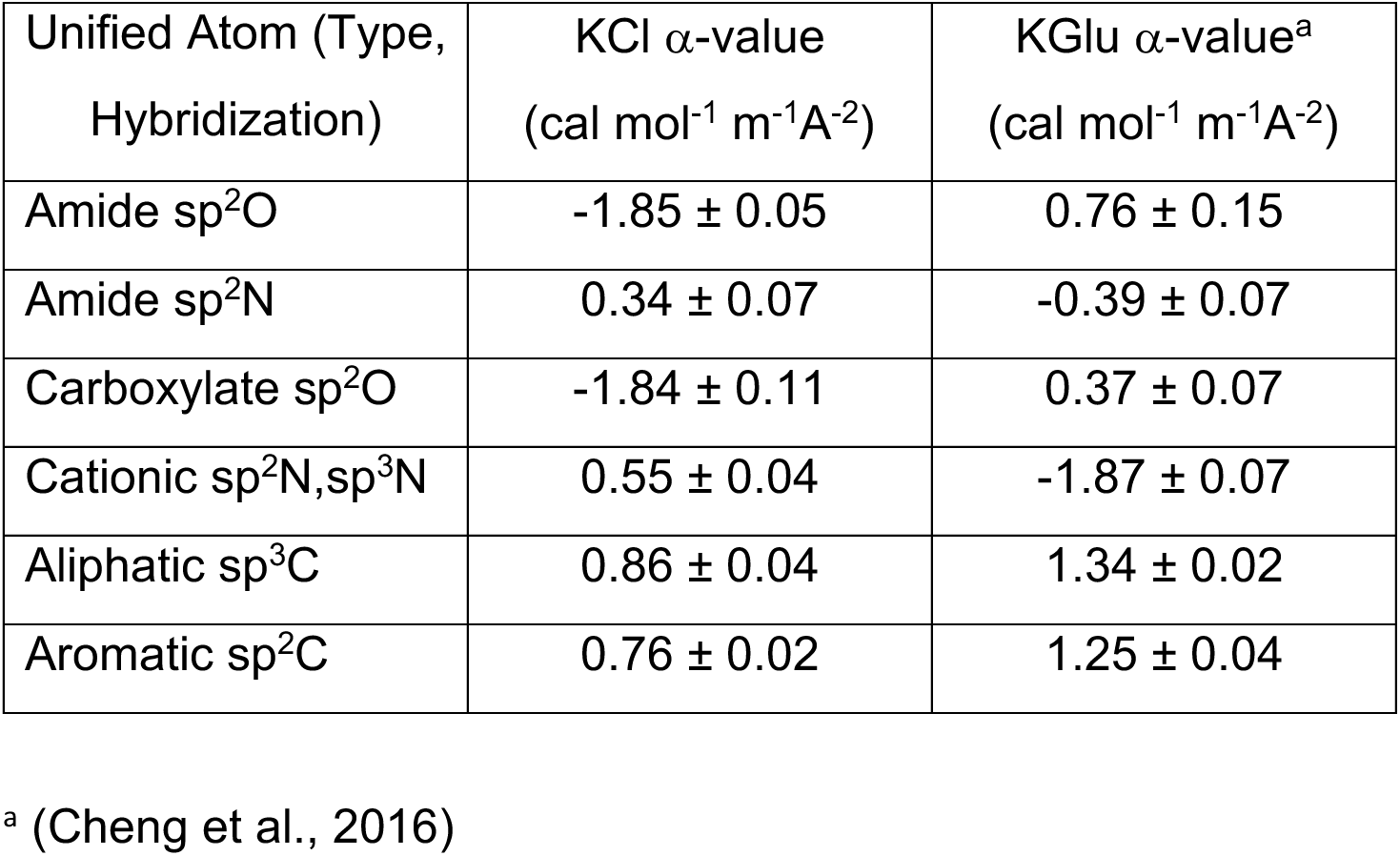
Intrinsic Strengths (*α*-Values) for Interactions of KCl with O, N, C Unified Atoms of Protein Model Compounds

**Table S3.** Contributions to Effects of KGlu and KCl on T_PS_ From Interactions of these Salts with G Backbone and Q,N Side Chain Amide Groups

| Amide Residue<br>Contribution to PS | G backbone | Q, N side chain |
| --- | --- | --- |
| Amide O ASA ($\text{\AA}^2$) | 28 $\text{\AA}^2$ | 45 $\text{\AA}^2$ |
| Amide N ASA ($\text{\AA}^2$) | 17 $\text{\AA}^2$ | 53 $\text{\AA}^2$ |
| KCl-Amide O<br>Contribution to PS | +52 cal mol<sup>-1</sup> m<sup>-1</sup> | +83 cal mol<sup>-1</sup> m<sup>-1</sup> |
| KCl-Amide N<br>Contribution to PS | -6 cal mol<sup>-1</sup> m<sup>-1</sup> | -18 cal mol<sup>-1</sup> m<sup>-1</sup> |
| Net KCl-Amide O,N<br>Contribution to PS | +46 cal mol<sup>-1</sup> m<sup>-1</sup> | +65 cal mol<sup>-1</sup> m<sup>-1</sup> |
| K-Glu Amide O<br>Contribution to PS | -21 cal mol<sup>-1</sup> m<sup>-1</sup> | -34 cal mol<sup>-1</sup> m<sup>-1</sup> |
| KGlu-Amide N<br>Contribution to PS | +7 cal mol<sup>-1</sup> m<sup>-1</sup> | +21 cal mol<sup>-1</sup> m<sup>-1</sup> |
| Net KGlu-Amide O,N<br>Contribution to PS | -14 cal mol<sup>-1</sup> m<sup>-1</sup> | -13 cal mol<sup>-1</sup> m<sup>-1</sup> |
| Total Possible KCl<br>Contribution per SSB<br>tetramer<sup>a</sup> | + 3.1 kcal mol<sup>-1</sup> m<sup>-1</sup> | +3.6 kcal mol<sup>-1</sup> m<sup>-1</sup> |
| Total Possible KGlu<br>Contribution per SSB<br>tetramer<sup>a</sup> | -1.0 kcal mol<sup>-1</sup> m<sup>-1</sup> | -0.7 kcal mol<sup>-1</sup> m<sup>-1</sup> |

<sup>a</sup> From salt interactions with 17x4 G amides and 14x4 Q,N side-chain amides.

## References

Alberti, S., Gladfelter, A. & Mittag, T. 2019. Considerations and Challenges in Studying Liquid-Liquid Phase Separation and Biomolecular Condensates. Cell, 176, 419–434.

Amiram, M., Quiroz, F. G., Callahan, D. J. & Chilkoti, A. 2011. A highly parallel method for synthesizing DNA repeats enables the discovery of ’smart’ protein polymers. Nat Mater, 10, 141–8.

Antony, E., Kozlov, A. G., Nguyen, B. & Lohman, T. M. 2012. Plasmodium falciparum SSB tetramer binds single-stranded DNA only in a fully wrapped mode. J Mol Biol, 420, 284–95.

Antony, E., Weiland, E., Yuan, Q., Manhart, C. M., Nguyen, B., Kozlov, G., Mchenry, C. S. & Lohman, T. M. 2013. Multiple C-Terminal Tails within a Single E. coli SSB Homotetramer Coordinate DNA Replication and Repair. J Mol Biol, 425, 4802–19.

Banani, S. F., Lee, H. O., Hyman, A. A. & Rosen, M. K. 2017. Biomolecular condensates: organizers of cellular biochemistry. Nat Rev Mol Cell Biol, 18, 285–298.

Bell, J. C., Liu, B. & Kowalczykowski, S. C. 2015. Imaging and energetics of single SSB-ssDNA molecules reveal intramolecular condensation and insight into RecOR function. Elife, 4, e08646.

Berkowitz, S. A. & Day, L. A. 1974. Molecular weight of single-stranded fd bacteriophage DNA. High speed equilibrium sedimentation and light scattering measurements. Biochemistry, 13, 4825–31.

Boeynaems, S., Bogaert, E., Kovacs, D., Konijnenberg, A., Timmerman, E., Volkov, A., Guharoy, M., DE Decker, M., Jaspers, T., Ryan, V. H., Janke, A. M., Baatsen, P., Vercruysse, T., Kolaitis, R. M., Daelemans, D., Taylor, J. P., Kedersha, N., Anderson, P., Impens, F., Sobott, F., Schymkowitz, J., Rousseau, F., Fawzi, N. L., Robberecht, W., VAN Damme, P., Tompa, P. & Van Den Bosch, L. 2017. Phase Separation of C9orf72 Dipeptide Repeats Perturbs Stress Granule Dynamics. Mol Cell, 65, 1044–1055 e5.

Boyko, S., Qi, X., Chen, T. H., Surewicz, K. & Surewicz, W. K. 2019. Liquid-liquid phase separation of tau protein: The crucial role of electrostatic interactions. J Biol Chem, 294, 11054–11059.

Brady, J. P., Farber, P. J., Sekhar, A., Lin, Y. H., Huang, R., Bah, A., Nott, T. J., Chan, H. S., Baldwin, A. J., Forman-Kay, J. D. & Kay, L. E. 2017. Structural and hydrodynamic properties of an intrinsically disordered region of a germ cell-specific protein on phase separation. Proc Natl Acad Sci U S A, 114, E8194–E8203.

Bujalowski, W. & Lohman, T. M. 1986. Escherichia coli single-strand binding protein forms multiple, distinct complexes with single-stranded DNA. Biochemistry, 25, 7799–802.

Bujalowski, W. & Lohman, T. M. 1987. Limited co-operativity in protein-nucleic acid interactions. A thermodynamic model for the interactions of Escherichia coli single strand binding protein with single-stranded nucleic acids in the "beaded", (SSB)65 mode. J Mol Biol, 195, 897–907.

Bujalowski, W., Overman, L. B. & Lohman, T. M. 1988. Binding mode transitions of Escherichia coli single strand binding protein-single-stranded DNA complexes. Cation, anion, pH, and binding density effects. J Biol Chem, 263, 4629–40.

Burke, K. A., Janke, A. M., Rhine, C. L. & Fawzi, N. L. 2015. Residue-by-Residue View of In Vitro FUS Granules that Bind the C-Terminal Domain of RNA Polymerase II. Mol Cell, 60, 231–41.

Capp, M. W., Pegram, L. M., Saecker, R. M., Kratz, M., Riccardi, D., Wendorff, T., Cannon, J. G. & Record, M. T., JR. 2009. Interactions of the osmolyte glycine betaine with molecular surfaces in water: thermodynamics, structural interpretation, and prediction of m-values. Biochemistry, 48, 10372–9.

Cayley, S., Lewis, B. A., Guttman, H. J. & Record JR., M. T. 1991. Characterization of the cytoplasm of *Escherichia coli* K-12 as a function of external osmolarity. Implications for protein-DNA interactions in vivo. J.Mol.Biol., 222, 281–300.

Chase, J. W. & Williams, K. R. 1986. Single-stranded DNA binding proteins required for DNA replication. Annu.Rev.Biochem., 55, 103–136.

Cheng, X., Guinn, E. J., Buechel, E., Wong, R., Sengupta, R., Shkel, I. A. & Record, M. T., JR. 2016. Basis of Protein Stabilization by K Glutamate: Unfavorable Interactions with Carbon, Oxygen Groups. Biophys J, 111, 1854–1865.

Cheng, X., Shkel, I. A., O’connor, K., Henrich, J., Molzahn, C., Lambert, D. & Record, M. T., Jr. 2017. Experimental Atom-by-Atom Dissection of Amide-Amide and Amide-Hydrocarbon Interactions in H2O. J Am Chem Soc, 139, 9885–9894.

Cheng, X., Shkel, I. A., O’connor, K. & Record, M. T., JR. 2020. Experimentally determined strengths of favorable and unfavorable interactions of amide atoms involved in protein self-assembly in water. Proc Natl Acad Sci U S A, 117, 27339–27345.

Choi, J. M., Holehouse, A. S. & Pappu, R. V. 2020. Physical Principles Underlying the Complex Biology of Intracellular Phase Transitions. Annu Rev Biophys, 49, 107–133.

Chong, P. A., Vernon, R. M. & Forman-Kay, J. D. 2018. RGG/RG Motif Regions in RNA Binding and Phase Separation. J Mol Biol, 430, 4650–4665.

Chrysogelos, S. & Griffith, J. 1982. Escherichia coli single-strand binding protein organizes single-stranded DNA in nucleosome-like units. Proc Natl Acad Sci U S A, 79, 5803–7.

Cinar, H., Fetahaj, Z., Cinar, S., Vernon, R. M., Chan, H. S. & Winter, R. H. A. 2019. Temperature, Hydrostatic Pressure, and Osmolyte Effects on Liquid-Liquid Phase Separation in Protein Condensates: Physical Chemistry and Biological Implications. Chemistry-a European Journal, 25, 13049–13069.

Cohan, M. C., Kyung SHINN, M., Lalmansingh, J. M. & Pappu, R. V. 2021. Uncovering non-random binary patterns within sequences of intrinsically disordered proteins. J Mol Biol, 167373.

Cohan, M. C. & Pappu, R. V. 2020. Making the Case for Disordered Proteins and Biomolecular Condensates in Bacteria. Trends Biochem Sci, 45, 668–680.

Dam, J. & Schuck, P. 2004. Calculating sedimentation coefficient distributions by direct modeling of sedimentation velocity concentration profiles. Methods Enzymol, 384, 185–212.

Deredge, D. J., Baker, J. T., Datta, K. & Licata, V. J. 2010. The glutamate effect on DNA binding by pol I DNA polymerases: osmotic stress and the effective reversal of salt linkage. J Mol Biol, 401, 223–38.

Dubiel, K., Myers, A. R., Kozlov, A. G., Yang, O., Zhang, J., Ha, T., Lohman, T. M. & Keck, J. L. 2019. Structural Mechanisms of Cooperative DNA Binding by Bacterial Single-Stranded DNA-Binding Proteins. J Mol Biol, 431, 178–195.

Falahati, H. & Haji-Akbari, A. 2019. Thermodynamically driven assemblies and liquid-liquid phase separations in biology. Soft Matter, 15, 1135–1154.

Ferrari, M. E., Bujalowski, W. & Lohman, T. M. 1994. Co-operative binding of Escherichia coli SSB tetramers to single-stranded DNA in the (SSB)35 binding mode. J Mol Biol, 236, 106–23.

Fetahaj, Z., Ostermeier, L., Cinar, H., Oliva, R. & Winter, R. 2021. Biomolecular Condensates under Extreme Martian Salt Conditions. J Am Chem Soc, 143, 5247–5259.

Genschel, J., Curth, U. & Urbanke, C. 2000. Interaction of E-coli single-stranded DNA binding protein (SSB) with exonuclease I. The carboxy-terminus of SSB is the recognition site for the nuclease. Biological Chemistry, 381, 183–192.

Griffith, J. D., Harris, L. D. & Register, J., 3RD 1984. Visualization of SSB-ssDNA complexes active in the assembly of stable RecA-DNA filaments. Cold Spring Harb Symp Quant Biol, 49, 553–9.

Hamon, L., Pastre, D., Dupaigne, P., Le Breton, C., Le Cam, E. & Pietrement, O. 2007. High-resolution AFM imaging of single-stranded DNA-binding (SSB) protein--DNA complexes. Nucleic Acids Res, 35, e58.

Harami, G. M., Kovacs, Z. J., Pancsa, R., Palinkas, J., Barath, V., Tarnok, K., Malnasi-Csizmadia, A. & Kovacs, M. 2020. Phase separation by ssDNA binding protein controlled via protein-protein and protein-DNA interactions. Proceedings of the National Academy of Sciences of the United States of America, 117, 26206–26217.

Hatch, K., Danilowicz, C., Coljee, V. & Prentiss, M. 2008. Measurement of the salt-dependent stabilization of partially open DNA by Escherichia coli SSB protein. Nucleic Acids Res, 36, 294–9.

Hofmeister, F. 1888. On the understanding of the effect of salts. Second report. On regularities in the precipitating effect od salts and their relationship to their physiological behavior. Naunyn-Schmiedebergs archive fuer Experimentelle Pathologie und Pharmakologie (Leipzig), 24, 247–260.

Holehouse, A. S. & Pappu, R. V. 2018. Functional Implications of Intracellular Phase Transitions. Biochemistry, 57, 2415–2423.

Kar, M., Posey, A. E., Dar, F., Hyman, A. A. & Pappu, R. V. 2021. Glycine-Rich Peptides from FUS Have an Intrinsic Ability to Self-Assemble into Fibers and Networked Fibrils. Biochemistry, 60, 3213–3222.

Knowles, D. B., Shkel, I. A., Phan, N. M., Sternke, M., Lingeman, E., Cheng, X., Cheng, L., O’connor, K. & Record, M. T. 2015. Chemical Interactions of Polyethylene Glycols (PEGs) and Glycerol with Protein Functional Groups: Applications to Effects of PEG and Glycerol on Protein Processes. Biochemistry, 54, 3528–42.

Kontur, W. S., Capp, M. W., Gries, T. J., Saecker, R. M. & Record, M. T., JR. 2010. Probing DNA binding, DNA opening, and assembly of a downstream clamp/jaw in Escherichia coli RNA polymerase-lambdaP(R) promoter complexes using salt and the physiological anion glutamate. Biochemistry, 49, 4361–73.

Kozlov, A. G., Cox, M. M. & Lohman, T. M. 2010a. Regulation of single-stranded DNA binding by the C termini of Escherichia coli single-stranded DNA-binding (SSB) protein. J Biol Chem, 285, 17246–52.

Kozlov, A. G., Jezewska, M. J., Bujalowski, W. & Lohman, T. M. 2010b. Binding specificity of Escherichia coli single-stranded DNA binding protein for the chi subunit of DNA pol III holoenzyme and PriA helicase. Biochemistry, 49, 3555–66.

Kozlov, A. G. & Lohman, T. M. 1999. Adenine base unstacking dominates the observed enthalpy and heat capacity changes for the Escherichia coli SSB tetramer binding to single-stranded oligoadenylates. Biochemistry, 38, 7388–7397.

Kozlov, A. G. & Lohman, T. M. 2002. Kinetic mechanism of direct transfer of Escherichia coli SSB tetramers between single-stranded DNA molecules. Biochemistry, 41, 11611–27.

Kozlov, A. G., Shinn, M. K. & Lohman, T. M. 2019. Regulation of Nearest-Neighbor Cooperative Binding of E. coli SSB Protein to DNA. Biophys J, 117, 2120–2140.

Kozlov, A. G., Shinn, M. K., Weiland, E. A. & Lohman, T. M. 2017. Glutamate promotes SSB protein-protein Interactions via intrinsically disordered regions. J Mol Biol, 429, 2790–2801.

Kozlov, A. G., Weiland, E., Mittal, A., Waldman, V., Antony, E., Fazio, N., Pappu, R. V. & Lohman, T. M. 2015. Intrinsically Disordered C-terminal Tails of *E. coli* Single Stranded DNA Binding Protein Regulate Cooperative Binding to Single Stranded DNA. Journal of Molecular Biology, 427, 763–774.

Krainer, G., Welsh, T. J., Joseph, J. A., Espinosa, J. R., Wittmann, S., De Csillery, E., Sridhar, A., Toprakcioglu, Z., Gudiskyte, G., Czekalska, M. A., Arter, W. E., Guillen-Boixet, J., Franzmann, T. M., Qamar, S., George-Hyslop, P. S., Hyman, A. A., Collepardo- Guevara, R., Alberti, S. & Knowles, T. P. J. 2021. Reentrant liquid condensate phase of proteins is stabilized by hydrophobic and non-ionic interactions. Nat Commun, 12, 1085.

Lee, K. S., Marciel, A. B., Kozlov, A. G., Schroeder, C. M., Lohman, T. M. & Ha, T. 2014. Ultrafast Redistribution of E. coli SSB along Long Single-Stranded DNA via Intersegment Transfer. J Mol Biol, 426, 2413–21.

Leirmo, S., Harrison, C., Cayley, D. S., Burgess, R. R. & Record, M. T., Jr. 1987. Replacement of potassium chloride by potassium glutamate dramatically enhances protein-DNA interactions in vitro. Biochemistry, 26, 2095–101.

Lohman, T. M. & Ferrari, M. E. 1994. Escherichia coli single-stranded DNA-binding protein: multiple DNA-binding modes and cooperativities. Annu Rev Biochem, 63, 527–70.

Lohman, T. M. & Overman, L. B. 1985. Two binding modes in Escherichia coli single strand binding protein-single stranded DNA complexes. Modulation by NaCl concentration. J Biol Chem, 260, 3594–603.

Lohman, T. M., Overman, L. B. & Datta, S. 1986. Salt-dependent changes in the DNA binding co-operativity of Escherichia coli single strand binding protein. J Mol Biol, 187, 603–15.

Lyons, D. F., Le, V., Bidwell, G. L., 3RD, Kramer, W. H., Lewis, E. A., Raucher, D. & Correia, J. J. 2013. Structural and hydrodynamic analysis of a novel drug delivery vector: ELP[V5G3A2-150]. Biophys J, 104, 2009–21.

Marceau, A. H., Bahng, S., Massoni, S. C., George, N. P., Sandler, S. J., Marians, K. J. & Keck, J. L. 2011. Structure of the SSB-DNA polymerase III interface and its role in DNA replication. EMBO J, 30, 4236–47.

Martin, E. W., Holehouse, A. S., Peran, I., Farag, M., Incicco, J. J., Bremer, A., Grace, C. R., Soranno, A., Pappu, R. V. & Mittag, T. 2020. Valence and patterning of aromatic residues determine the phase behavior of prion-like domains. Science, 367, 694–699.

Mason, B. D., Zhang-Van Enk, J., Zhang, L., Remmele, R. L. & Zhang, J. 2010. Liquid-Liquid Phase Separation of a Monoclonal Antibody and Nonmonotonic Influence of Hofmeister Anions. Biophysical Journal, 99, 3792–3800.

Meyer, R. R. & Laine, P. S. 1990. The single-stranded DNA-binding protein of *Escherichia coli*. Microbiological Reviews, 54, 342–380.

Murthy, A. C., Tang, W. S., Jovic, N., Janke, A. M., Seo, D. H., Perdikari, T. M., Mittal, J. & Fawzi, N. L. 2021. Molecular interactions contributing to FUS SYGQ LC-RGG phase separation and co-partitioning with RNA polymerase II heptads. Nat Struct Mol Biol, 28, 923–935.

Nott, T. J., Petsalaki, E., Farber, P., Jervis, D., Fussner, E., Plochowietz, A., Craggs, T. D., Bazett-Jones, D. P., Pawson, T., Forman-Kay, J. D. & Baldwin, A. J. 2015. Phase transition of a disordered nuage protein generates environmentally responsive membraneless organelles. Mol Cell, 57, 936–947.

Overman, L. B., Bujalowski, W. & Lohman, T. M. 1988. Equilibrium binding of Escherichia coli single-strand binding protein to single-stranded nucleic acids in the (SSB)65 binding mode. Cation and anion effects and polynucleotide specificity. Biochemistry, 27, 456–71.

Pegram, L. M. & Record, M. T., JR. 2007. Hofmeister salt effects on surface tension arise from partitioning of anions and cations between bulk water and the air-water interface. J Phys Chem B, 111, 5411–7.

Petsev, D. N., Wu, X. X., Galkin, O. & Vekilov, P. G. 2003. Thermodynamic functions of concentrated protein solutions from phase equilibria. Journal of Physical Chemistry B, 107, 3921–3926.

Posey, A. E., Holehouse, A. S. & Pappu, R. V. 2018. Phase Separation of Intrinsically Disordered Proteins. Intrinsically Disordered Proteins, 611, 1–30.

Raghunathan, S., Kozlov, A. G., Lohman, T. M. & Waksman, G. 2000. Structure of the DNA binding domain of E-coli SSB bound to ssDNA. Nature Structural Biology, 7, 648–652.

Record JR., M. T., Courtenay, E. S., Cayley, D. S. & Guttman, H. J. 1998. Responses of *E. coli* to osmotic stress: large changes in amounts of cytoplasmic solutes and water. TIBS, 23, 143–148.

Richey, B., Cayley, D. S., Mossing, M. C., Kolka, C., Anderson, C. F., Farrar, T. C. & Record, M. T., JR. 1987. Variability of the intracellular ionic environment of Escherichia coli. Differences between in vitro and in vivo effects of ion concentrations on protein-DNA interactions and gene expression. J Biol Chem, 262, 7157–64.

Robinson, R. A. & Stokes, R. H. 1961. Activity coefficients in aqueous solutions of sucrose, mannitol and their mixtures at 25°C. J Phys Chem, 65, 1954–1958.

Roy, R., Kozlov, A. G., Lohman, T. M. & Ha, T. 2007. Dynamic Structural Rearrangements Between DNA Binding Modes of E. coli SSB Protein. J Mol Biol, 369, 1244–57.

Roy, R., Kozlov, A. G., Lohman, T. M. & Ha, T. 2009. SSB protein diffusion on single-stranded DNA stimulates RecA filament formation. Nature, 461, 1092–7.

Rubinstein, M., & Colby, R.H. 2003. Polymer physics: Chapter 4, New York, NY, Oxford University Press.

Ryzhikov, M. & Korolev, S. 2012. Structural studies of SSB interaction with RecO. Methods Mol Biol, 922, 123–31.

Schaub, J. M., Zhang, H., Soniat, M. M. & Finkelstein, I. J. 2018. Assessing Protein Dynamics on Low-Complexity Single-Stranded DNA Curtains. Langmuir, 34, 14882–14890.

Sengupta, R., Capp, M. W., Shkel, I. A. & Record, M. T., JR. 2017. The mechanism and high-free-energy transition state of lac repressor-lac operator interaction. Nucleic Acids Res, 45, 12671–12680.

Sengupta, R., Pantel, A., Cheng, X., Shkel, I., Peran, I., Stenzoski, N., Raleigh, D. P. & Record, M. T., Jr. 2016. Positioning the Intracellular Salt Potassium Glutamate in the Hofmeister Series by Chemical Unfolding Studies of NTL9. Biochemistry, 55, 2251–9.

Shereda, R. D., Kozlov, A. G., Lohman, T. M., Cox, M. M. & Keck, J. L. 2008. SSB as an organizer/mobilizer of genome maintenance complexes. Crit Rev Biochem Mol Biol, 43, 289–318.

Shereda, R. D., Reiter, N. J., Butcher, S. E. & Keck, J. L. 2009. Identification of the SSB binding site on E. coli RecQ reveals a conserved surface for binding SSB’s C terminus. J Mol Biol, 386, 612–25.

Shin, Y. & Brangwynne, C. P. 2017. Liquid phase condensation in cell physiology and disease. Science, 357.

Shinn, M. K., Kozlov, A. G., Nguyen, B., Bujalowski, W. M. & Lohman, T. M. 2019. Are the intrinsically disordered linkers involved in SSB binding to accessory proteins? Nucleic Acids Res, 47, 8581–8594.

Sigal, N., Delius, H., Kornberg, T., Gefter, M. L. & Alberts, B. 1972. A DNA-unwinding protein isolated from Escherichia coli: its interaction with DNA and with DNA polymerases. Proc Natl Acad Sci U S A, 69, 3537–41.

Sokoloski, J. E., Kozlov, A. G., Galletto, R. & Lohman, T. M. 2016. Chemo-mechanical pushing of proteins along single-stranded DNA. Proc Natl Acad Sci U S A, 113, 6194–9.

Soniat, M. M., Myler, L. R., Schaub, J. M., Kim, Y., Gallardo, I. F. & Finkelstein, I. J. 2017. Next-Generation DNA Curtains for Single-Molecule Studies of Homologous Recombination. *DNA Repair Enzymes: Structure*, Biophysics, and Mechanism, 592, 259–281.

Spenkelink, L. M., Lewis, J. S., Jergic, S., Xu, Z. Q., Robinson, A., Dixon, N. E. & Van Oijen, A. M. 2019. Recycling of single-stranded DNA-binding protein by the bacterial replisome. Nucleic Acids Res.

Su, X. C., Wang, Y., Yagi, H., Shishmarev, D., Mason, C. E., Smith, P. J., Vandevenne, M., Dixon, N. E. & Otting, G. 2014. Bound or free: interaction of the C-terminal domain of Escherichia coli single-stranded DNA-binding protein (SSB) with the tetrameric core of SSB. Biochemistry, 53, 1925–34.

Suksombat, S., Khafizov, R., Kozlov, A. G., Lohman, T. M. & Chemla, Y. 2015. Structural dynamics of E. coli single-stranded DNA binding protein reveal DNA wrapping and unwrapping pathways. Elife, 4.

Taratuta, V. G., Holschbach, A., Thurston, G. M., Blankschtein, D. & Benedek, G. B. 1990. Liquid Liquid-Phase Separation of Aqueous Lysozyme Solutions - Effects of Ph and Salt Identity. Journal of Physical Chemistry, 94, 2140–2144.

Thomson, J. A., Schurtenberger, P., Thurston, G. M. & Benedek, G. B. 1987. Binary-Liquid Phase-Separation and Critical Phenomena in a Protein Water Solution. Proceedings of the National Academy of Sciences of the United States of America, 84, 7079–7083.

Tsang, B., Arsenault, J., Vernon, R. M., Lin, H., Sonenberg, N., Wang, L. Y., Bah, A. & Forman-Kay, J. D. 2019. Phosphoregulated FMRP phase separation models activity-dependent translation through bidirectional control of mRNA granule formation. Proc Natl Acad Sci U S A, 116, 4218–4227.

Van Der Lee, R., Buljan, M., Lang, B., Weatheritt, R. J., Daughdrill, G. W., Dunker, A. K., Fuxreiter, M., Gough, J., Gsponer, J., Jones, D. T., Kim, P. M., Kriwacki, R. W., Oldfield, C. J., Pappu, R. V., Tompa, P., Uversky, V. N., Wright, P. E. & Babu, M. M. 2014. Classification of Intrinsically Disordered Regions and Proteins. Chem Rev.

Van Holst, J., Kersten, S. R. A. & Hogendoorn, K. J. A. 2008. Physicochemical Properties of Several Aqueous Potassium Amino Acid Salts. J. Chem. Eng. Data, 53, 186–1291.

Von Hippel, P. H. & Schleich, T. 1969. Ion effects on the solution structure of biological macromolecules. Accts.Chem.Res., 2, 257–265.

Wold, M. S. 1997. Replication protein A: a heterotrimeric, single-stranded DNA-binding protein required for eukaryotic DNA metabolism. Annu Rev Biochem, 66, 61–92.

Yu, C., Tan, H. Y., Choi, M., Stanenas, A. J., Byrd, A. K., K, D. R., Cohan, C. & Bianco, P. R. 2016. SSB binds to the RecG and PriA helicases in vivo in the absence of DNA. Genes Cells, 21, 163–84.

Zhang, H. S., Schaub, J. M. & Finkelstein, I. J. 2020. RADX condenses single-stranded DNA to antagonize RAD51 loading. Nucleic Acids Research, 48, 7834–7843.

Zhang, Y. & Cremer, P. S. 2009. The inverse and direct Hofmeister series for lysozyme. Proc Natl Acad Sci U S A, 106, 15249–53.

Zhao, T., Liu, Y., Wang, Z., He, R., Xiang Zhang, J., Xu, F., Lei, M., Deci, M. B., Nguyen, J. & Bianco, P. R. 2019. Super-resolution imaging reveals changes in Escherichia coli SSB localization in response to DNA damage. Genes Cells, 24, 814–826.

Zhou, R., Kozlov, A. G., Roy, R., Zhang, J., Korolev, S., Lohman, T. M. & Ha, T. 2011. SSB functions as a sliding platform that migrates on DNA via reptation. Cell, 146, 222–32.

